# Biodiversity of the Genus *Trichoderma* in the Rhizosphere of Coffee (*Coffea arabica*) Plants in Ethiopia and their Potential Use in Biocontrol of Coffee Wilt Disease

**DOI:** 10.1101/2022.01.24.477504

**Authors:** Afrasa Mulatu, Negussie Megersa, Tariku Abena, Selvaraju Kanagarajan, Qinsong Liu, Tesfaye Alemu, Ramesh R. Vetukuri

## Abstract

The present study investigated the distribution status and biodiversity of *Trichoderma* species surveyed from coffee rhizosphere soil samples from Ethiopia and their potential for biocontrol of coffee wilt disease (CWD) caused by *Fusarium xylarioides*. *Trichoderma* isolates were identified based on molecular approaches and morphological characteristics followed by biodiversity analysis using different biodiversity indices. The antagonistic potential of *Trichoderma* isolates was evaluated against *F. xylarioides* using the dual confrontation technique and agar diffusion bioassays. A relatively high diversity of species was observed including 16 taxa and 11 undescribed isolates. *Trichoderma asperellum*, *T. asperelloides* and *T. longibrachiatum* were classified as abundant species, with dominance (Y) values of 0.062, 0.056 and 0.034, respectively. *Trichoderma asperellum* was the most abundant species (comprising 39.6% of all isolates) in all investigated coffee ecosystems. Shannon’s biodiversity index (H), the evenness (E), Simpson’s biodiversity index (D), and the abundance index (J) were calculated for each coffee ecosystem, revealing that species diversity and evenness were highest in the Jimma zone (H =1.97, E = 0.76, D = 0.91, J = 2.73). The average diversity values for *Trichoderma* species originating from the coffee ecosystem were H = 1.77, D = 0.7, E = 0.75 and J = 2.4. *In vitro* confrontation experiments revealed that *T. asperellum* AU131 and *T. longibrachiatum* AU158 reduced the mycelial growth of *F. xylarioides* by over 80%. The potential use of these *Trichoderma* species for disease management of *F. xylarioides* and to reduce its impact on coffee cultivation is discussed in relation to Ethiopia’s ongoing coffee wilt disease crisis.

## 1. Introduction

*Trichoderma* species are widely found in different soil types, ecosystems and climatic zones, and categorized based on their metabolic, physiological, and genetic diversity features [1]. They are economically significant because of their functions as primary decomposers, producers of antimicrobial compounds and enzymes, and their use as biocontrol agents against diverse phytopathogens [2–5]. Many research studies have revealed that in addition to directly inhibiting phytopathogens growth via mycoparasitism, antibiosis, and competition [6], some *Trichoderma* species have beneficial effects on plants resulting from plant growth promotion, solubilization of soil micro- and macro-nutrients [7], and activation of plant systemic resistance [8], in a multifaceted three-way interaction between antagonist, phytopathogens, and host plants [9]. To date, studies on *Trichoderma* diversity have mainly been conducted in Asia, Europe, and America [10]; there have been few investigations into the diversity and distribution of *Trichoderma* in Africa, with the exception of some studies targeting specific ecological niches [11,12]. In particular, there has been only one published study on *Trichoderma* species inhabiting coffee plants, which focused on species isolated from the rhizosphere of *C*. *arabica* in Ethiopia [13].

Morphological characterization and distinction was first used by Rifai [14] and later by Bisset [15–18] to investigate the diversity and evolution of *Trichoderma* species. However, species identification and delimination based on morphology alone is very difficult, making such approaches unreliable and subjective [19]. A more reliable approach is molecular phylogenetic analysis based on DNA sequencing data; over 375 *Trichoderma* species have been validly described and characterized in this way [20]. Reliable phylogenetic information is also important for studying the diversity of secondary metabolites of *Trichoderma* species. Consequently, molecular biological analysis is essential for the accurate identification of *Trichoderma* [21]. The internal transcribed spacer (ITS) is a widely used “universal” fungal molecular barcode [22,23]. However, it has low species resolution in the genus *Trichoderma* [24]. Therefore, the sequence of translation elongation factor 1-alpha (*TEF1-α*) was recommended as an alternative molecular barcode for phylogenetic analysis of this genus [24].

*Trichoderma* species stand out among rhizospheric microorganisms due to their high biocontrol potential and their ability to facilitate nutrient uptake by plants while also providing protection against phytopathogens [25]. To maximize their beneficial effects on crop plants, it is essential to evaluate the functional and structural diversity of *Trichoderma* species found in specific agro-climatic conditions. The rhizosphere of coffee exhibits particularly high diversity with a wide range of putative *Trichoderma* species and is a hotspot for the evolution of this genus [13]*. Trichoderma* species have been extensively studied and used as biocontrol agents against diverse plant pathogens including bacteria [26,27], fungi [28], oomycetes [29], and nematodes [30] for many different crops and agro-climatic conditions [31].

Ethiopia is the center of origin for Arabica coffee (*Coffea arabica* L.) and hosts a large germplasm diversity. It is also Africa’s largest coffee producer and the world’s fifth-largest coffee exporter, with a forecasted production of 457,200 metric tons (MT) in 2021/2022, having a value in excess of 900 million USD [32,33]. Coffee cultivation provides a livelihood for around 25 million [34,35], accounting for 25-30% of total export incomes [36]. In addition to the worldwide reputation of Ethiopia’s genetic resources, coffee plays a major role in the national economy and the livelihoods of approximately 4.5 million coffee farmers [37,38]. Despite its leading position in coffee cultivation in Africa, the Ethiopian coffee sector is underachieving due to the rise of various fungal and bacterial diseases, and these pressures are predicted to increase with climate change [39,40]. During the last decade of the 20th century, Coffee Wilt Disease (CWD) caused by *Fusarium xylarioides* become the principal production constraint for Arabica coffee in Ethiopia, Uganda, the Democratic Republic of Congo (DRC) and Tanzania [40]. The yearly coffee yield loss due to CWD in Ethiopia is estimated to be 30 - 40% [40–43]. CWD incidence is greatly affected by the farming system, with much higher rates in garden and plantation coffee. CWD has conventionally been managed by uprooting and burning the affected coffee plant and using resistant varieties [44].

The potential use of *Trichoderma* species for plant pathogen control is now well documented, although this approach is largely unexploited for many diseases of tropical perennial crops. Therefore, given the importance of coffee in Ethiopia’s national economy, the damaging nature of CWD, the limited availability of resistant crop lines, and the lack of information on the biocontrol of CWD, a study on the potential of *Trichoderma* species to suppress the growth of *F. xylarioides* is needed to identify new genomic resources for management of this pathogen. Screening the biodiversity of different coffee ecosystems and the ecophysiology of *Trichoderma* species from a genomic perspective and analyzing their diversity will provide important insights into the potential value of *Trichoderma* for controlling CWD in the future.

The prospect of influencing coffee rhizosphere by inoculating potential *Trichoderma* species to control CWD, enhance coffee growth and health was substantially studied under laboratory, greenhouse and field conditions (Mulatu A., unpublished data). However, the reduced efficiency of biocontrol agents under field condition is hindered due to their ability to adapt to local biotic and abiotic environmental conditions. To understand this phenomenon, it is necessary to study the geographical distribution and habitat preference of biocontrol agents in the rhizosphere. Hence, the present investigation was undertaken to study the distribution and biodiversity patterns of *Trichoderma* species in major coffee growing regions of Ethiopia with the long-term objective of assessing their potential as biocontrol agents of CWD.

## 2. Materials and Methods

### 2.1 Collection of Soil Samples and Isolation of Trichoderma species

*Trichoderma* isolates were collected from ten major Ethiopian coffee growing areas (Jimma, Kaffa, Benchi Maji, Sheka, Bunno Bedele, Bale, Sidama, Gedio, West Wollega and West Guji) in different agro-climatic zones. *Trichoderma* isolates were obtained from coffee rhizosphere soil gathered during surveys conducted between May 2016 and August 2017. The surveys covered all major coffee growing areas of Ethiopia’s southern, western, and southwestern regions. During soil collection, the upper surface soil litter (4–6 cm) was discarded, and 200 g soil samples were collected from a depth of approximately 10-15 cm. Over 184 soil samples were obtained from 28 districts (categorized under 10 zones) along the main roads (Figure 1 and Table S1). The soil samples were placed in sterile polyethylene bags, transported to the laboratory, and processed immediately. The strains were isolated using *Trichoderma* Specific Medium (TSM) according to previously reported methods Gil*, et al.* [45] and Saravanakumar*, et al.* [46] and purified by sub-culturing on potato dextrose agar (PDA). *Fusarium xylarioides* (DSM No. 62457, strain: IMB 11646), the causative agent of coffee wilt disease [9,47,48], was used as a test pathogen to evaluate the biocontrol potential of *Trichoderma* species.

**Figure 1.**
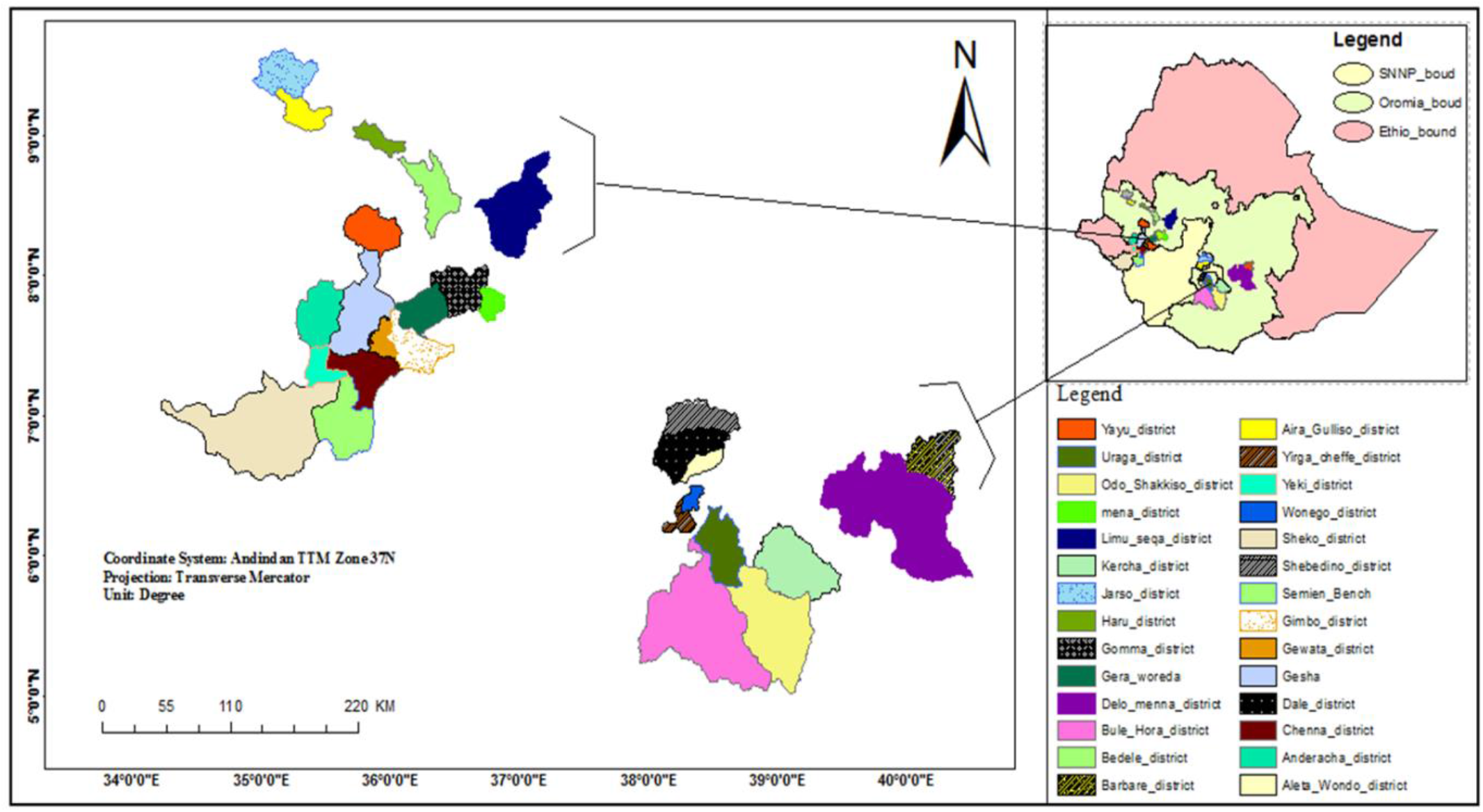
Map of study areas and illustration of the geographical locations of districts from which rhizospheric soil samples were collected, Ethiopia. SNNP = South Nations and Nationalities Peoples region.

### 2.2 Morphological Characteristics

The *Trichoderma* isolates were characterized based on their morphology by growing them on PDA at 28 ± 2°C for 5 days following the protocol described by Samuels and Hebbar [49]. The *Trichoderma* colonies were visually observed to determine their color (obverse and reverse), texture, margin, and sporulation. All *Trichoderma* isolates were classified and identified at the species level using morphological characteristics as suggested by Rifai [31] and Leahy and Colwell [50]. For further experiments and long-term storage, *Trichoderma* isolates were sub-cultured and slants were prepared in cryovials overlaid with 20 % glycerol and stored at −80 °C.

### 2.3 DNA Extraction, PCR Amplification and Sequencing

Genomic DNA was extracted according to Gontia-Mishra et al. (2014). Polymerase chain reaction (PCR) amplification of the *TEF1-α* region was performed using EF2-EF1728M primer following the conditions given by White*, et al.* [51]. PCR amplifications were carried out in a total reaction volume of 12.5 μl, including 0.25 μl of each primer, 1.25 μl of BSA, 6.25 of Taq polymerase [including dNTPs], 0.25 μl of genomic DNA [30 ng/μl]; 0.25 μl DMSO and 4 μl of sterile ultrapure water. PCR conditions for *TEF1-α*, conditions were 94 °C/2 min, followed by nine cycles at 94 °C/35 s, 66 °C/ 55 s, and 35 cycles at 94 °C/35 s, 56 °C/55 s and 72 °C/1 min 30 s. PCR products were visualized by Gelred (Thermo Fisher Scientific, Germany) staining following electrophoresis of 4 μl of each product in 1% agarose gel. The PCR products were sequenced by the Eurofins Sanger sequencing facility, Germany.

### 2.4 Phylogenetic Analysis

Consensus sequences were assembled from forward and reverse sequencing chromatograms using the CLC Main Workbench 8.1 software packages. *Tef1-α* contigs of all isolates were compared to homologous sequences deposited in the NCBI GenBank database. Sequences generated and used in the current study were deposited in this database (Table 1). Sequences utilized from other studies were retrieved from the NCBI GenBank database for use in our phylogenetic analyses. Sequence alignments were carried out using MUSCLE as implemented in MEGA 10 [52]. Before phylogenetic analyses, the most appropriate nucleotide substitution model for each locus was chosen using MRMODELTEST v.2126. The nucleotide substitution model for *TEF1-α* was HKY + I + G. The evolutionary history or consensus tree was inferred using the Maximum Likelihood test [53]. *Trichoderma* species matching with the isolates obtained in this work were retrieved and used to construct the phylogenetic tree, including two *Nectria* species as the outgroup. Nodal robustness was checked using the bootstrap method and phylogenetic robustness was evaluated using 1000 replicates. Only sequences that matched published results identified through BLASTN searches with >97% sequence identity and an e-value of zero were used.

**Table 1.**
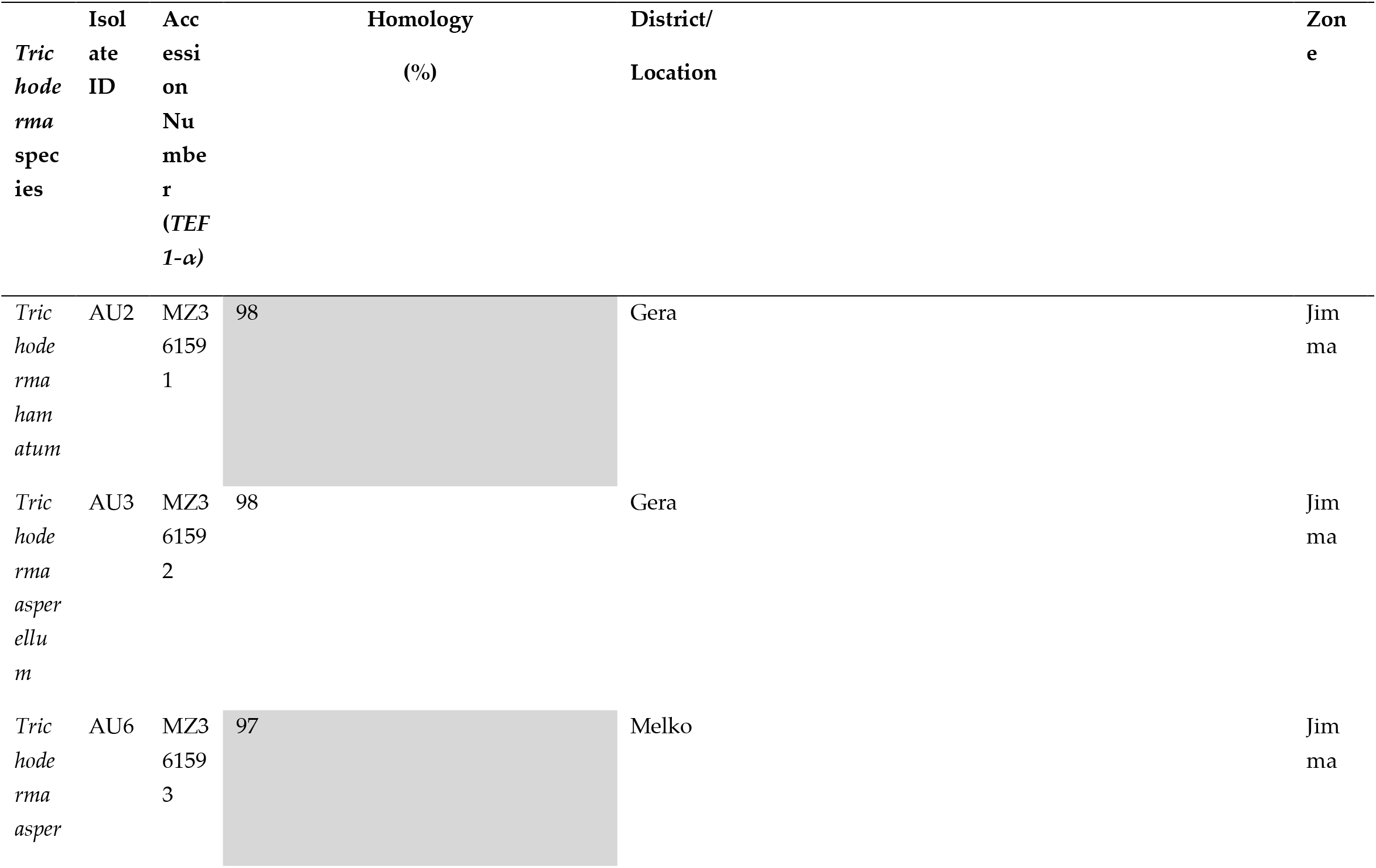
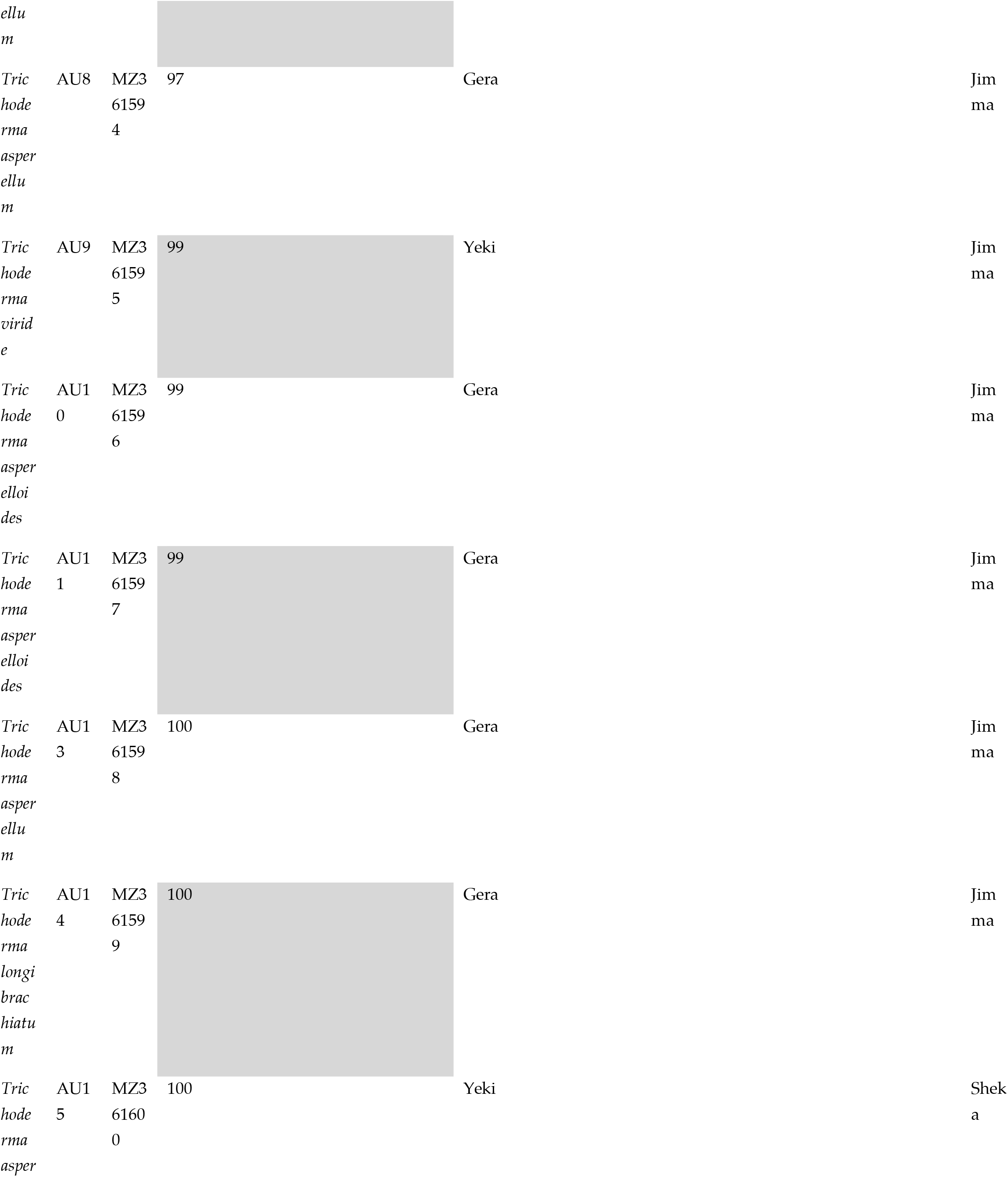
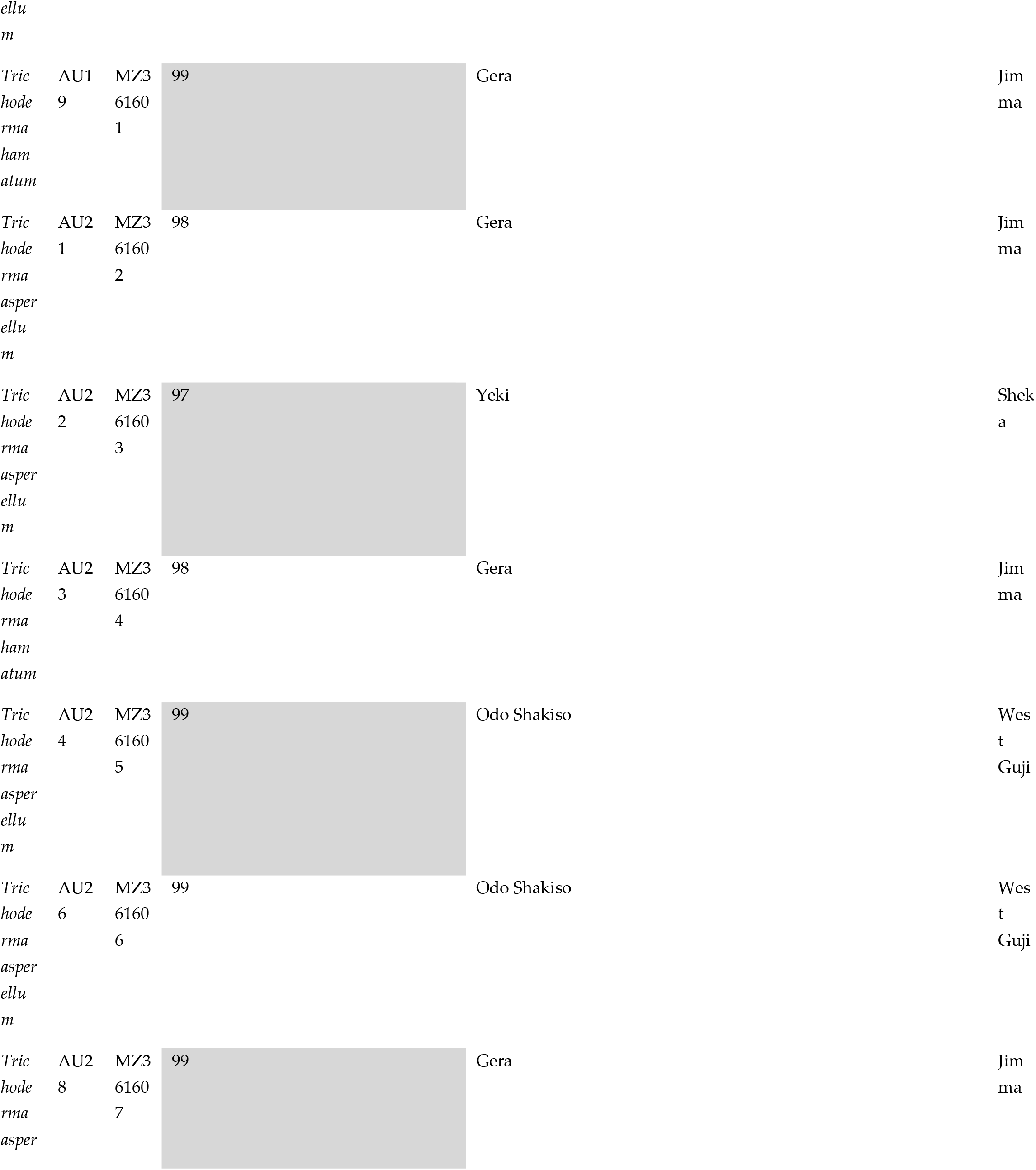
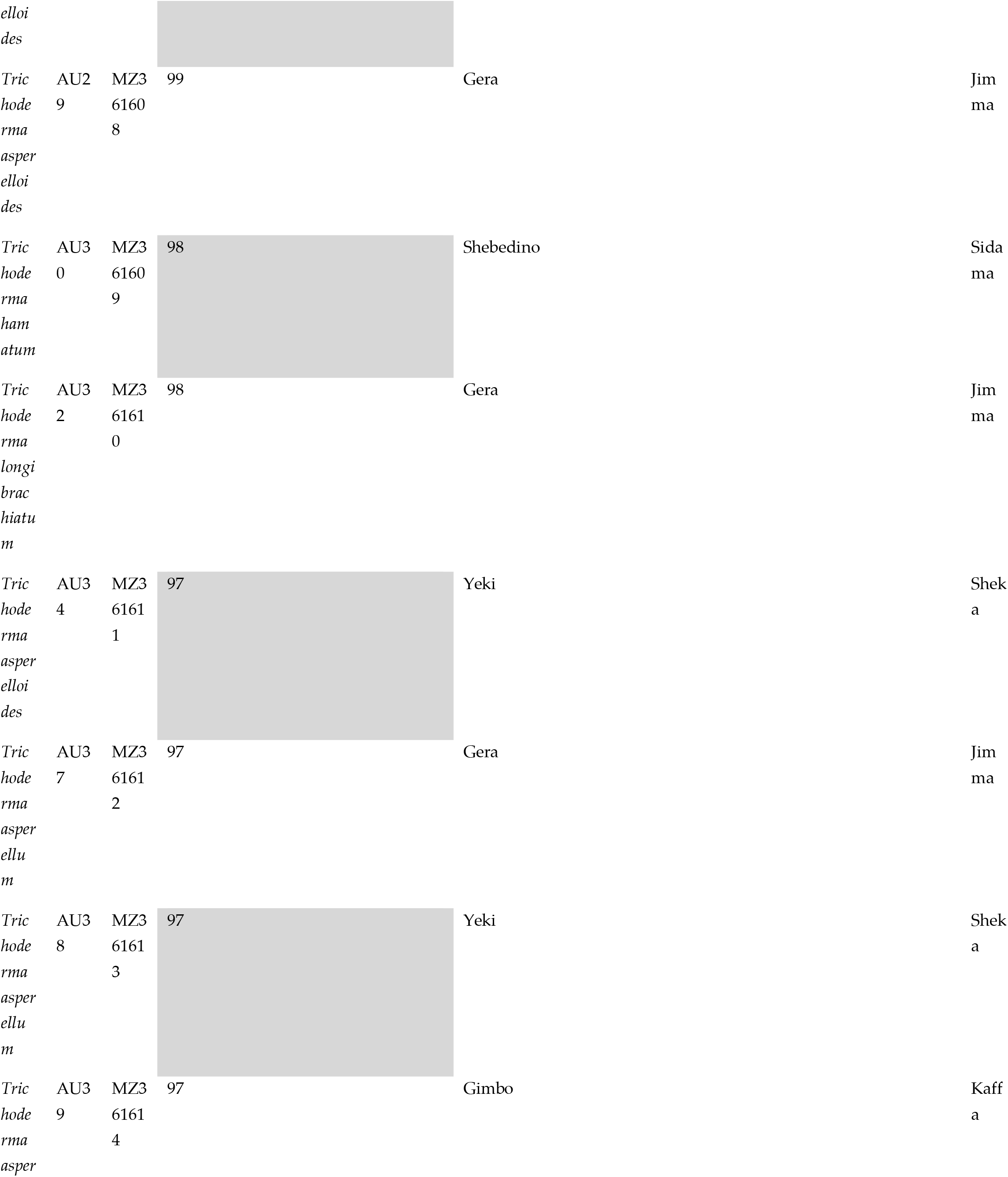
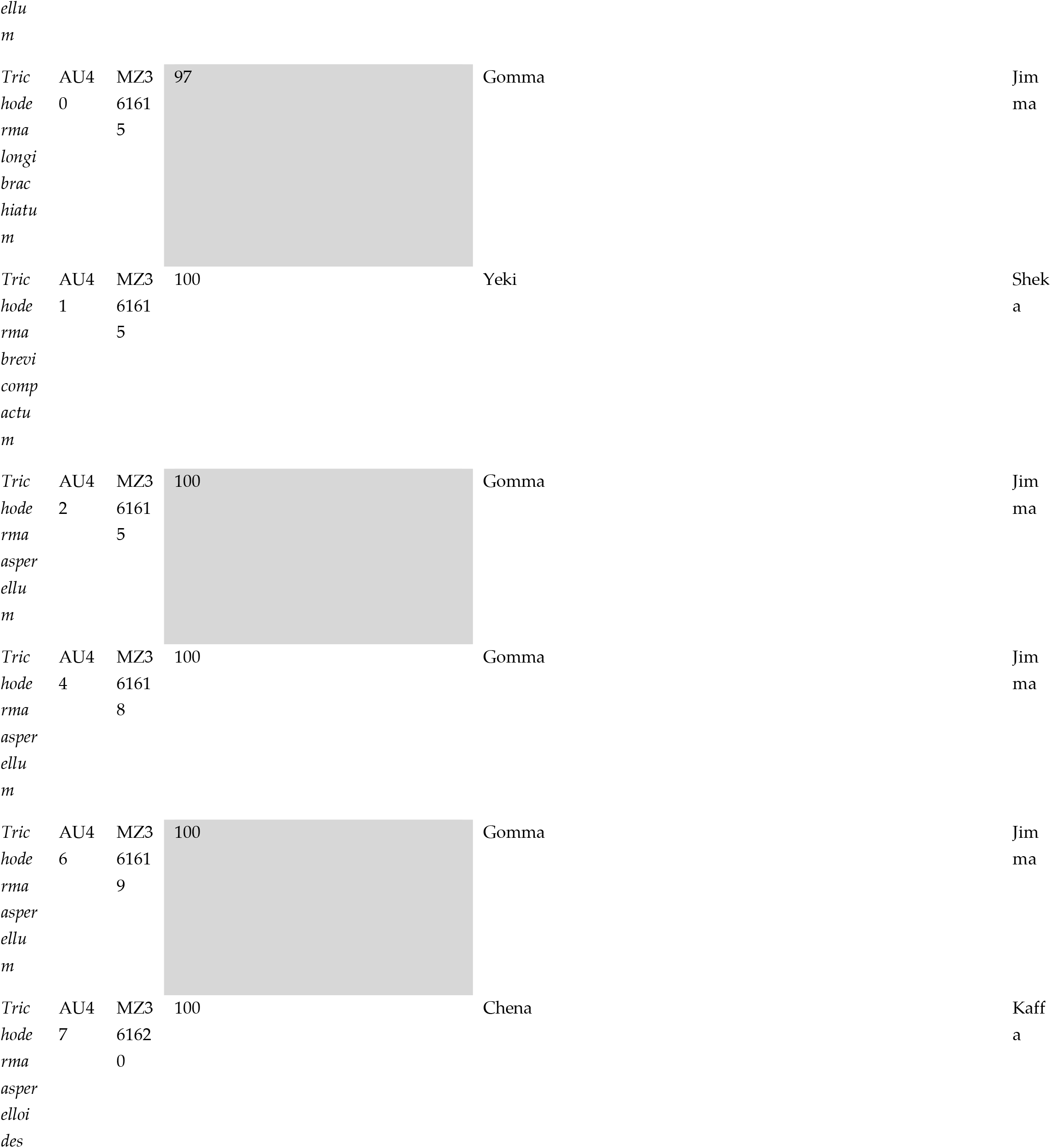
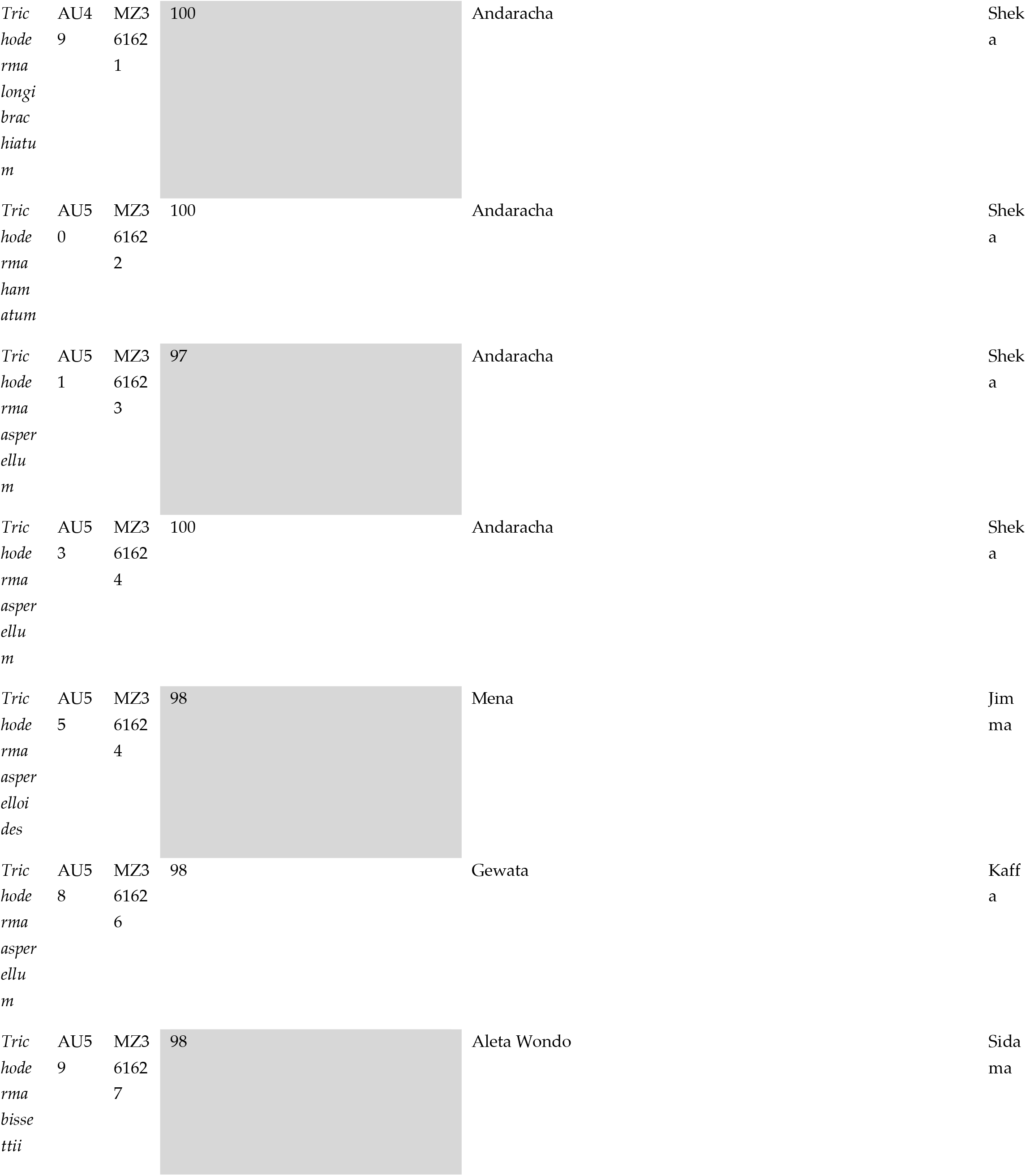
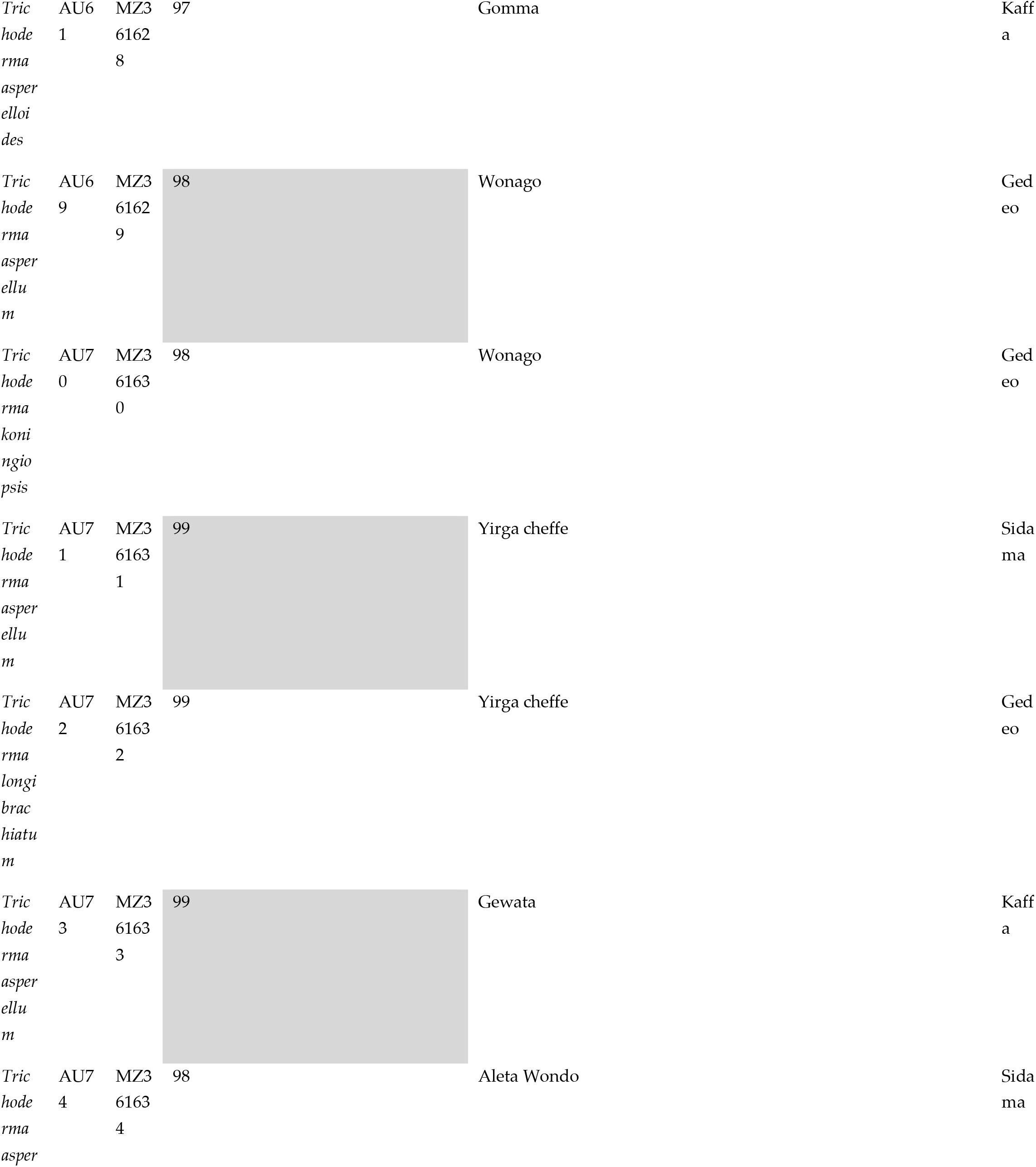
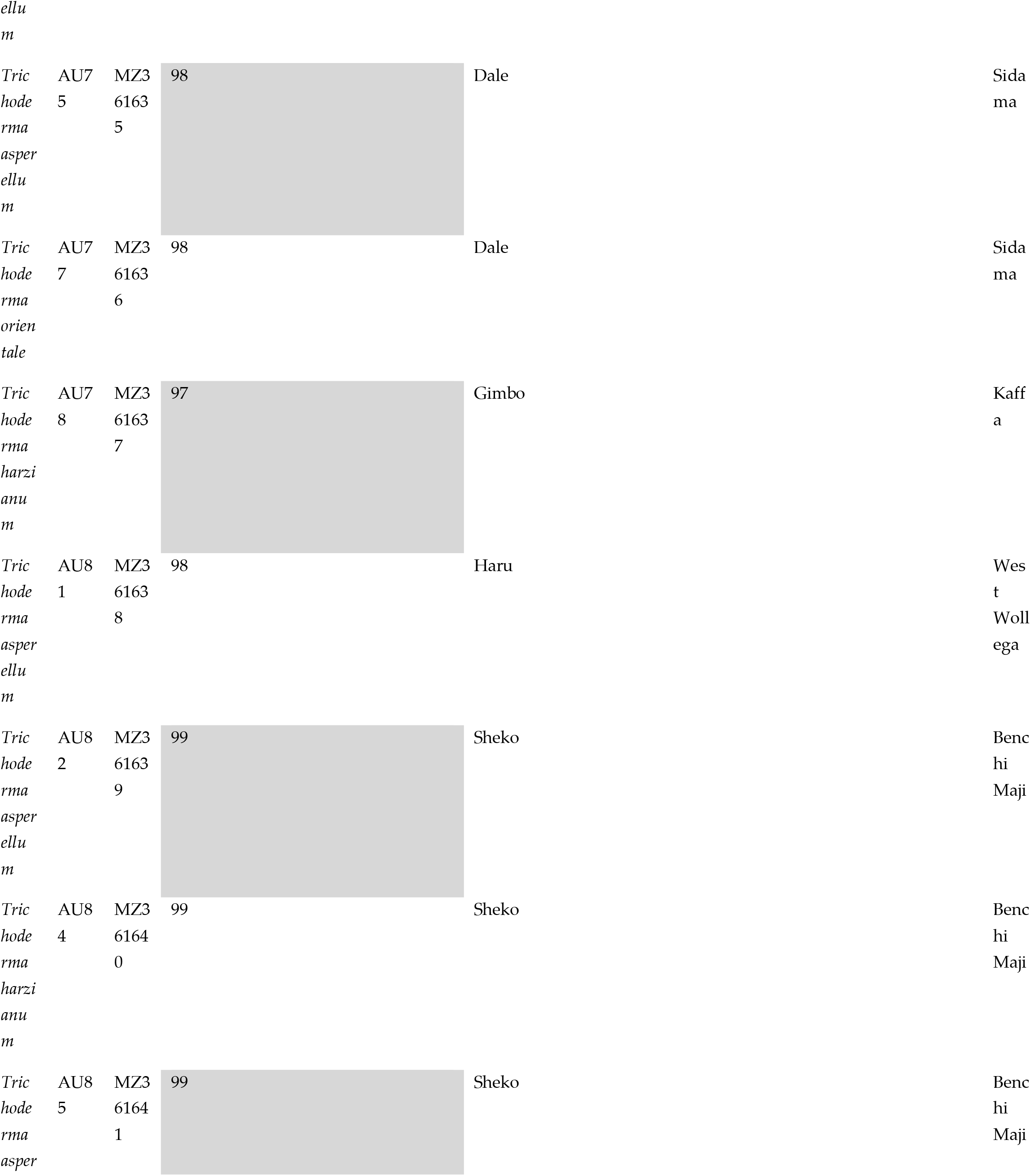
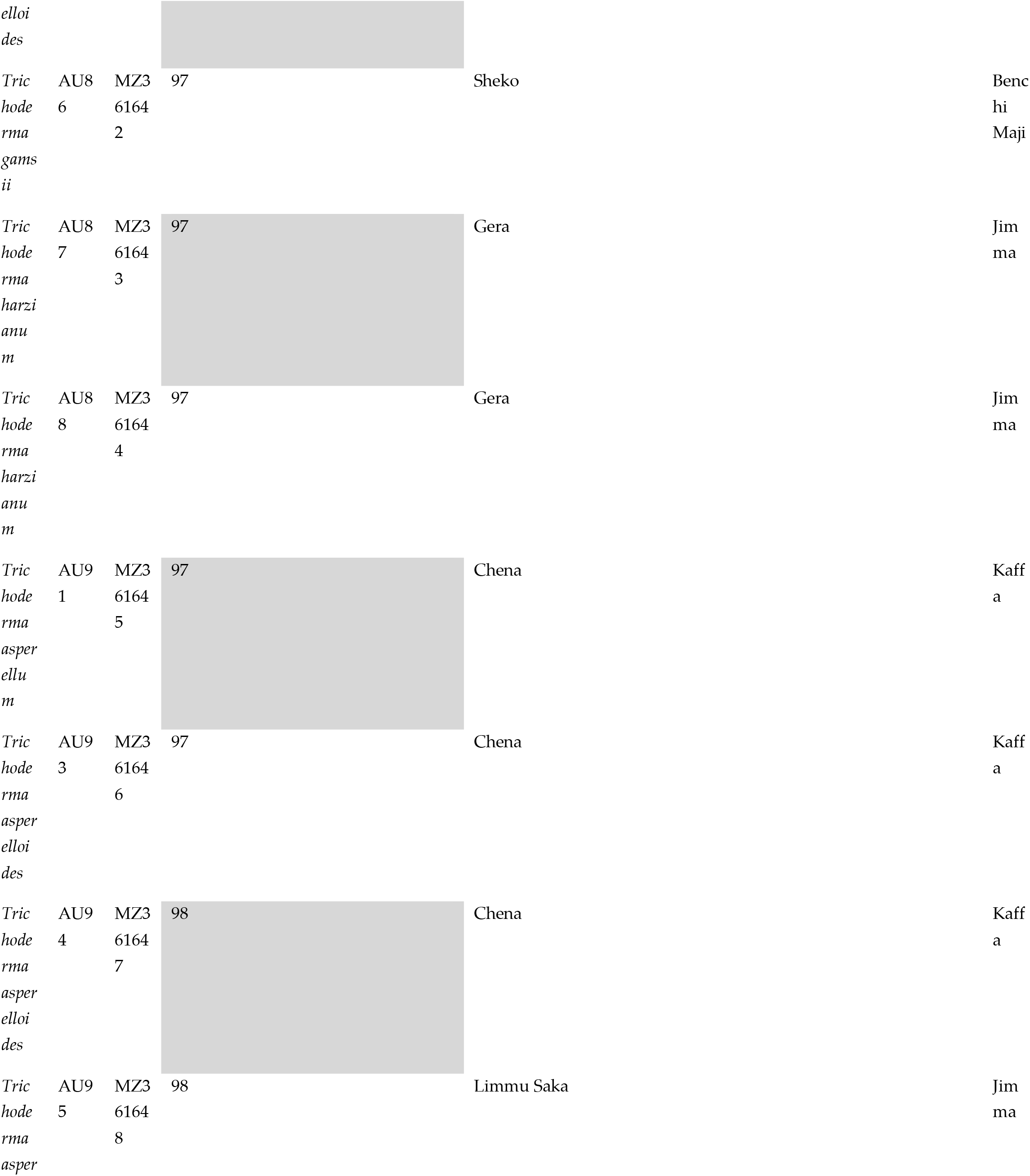
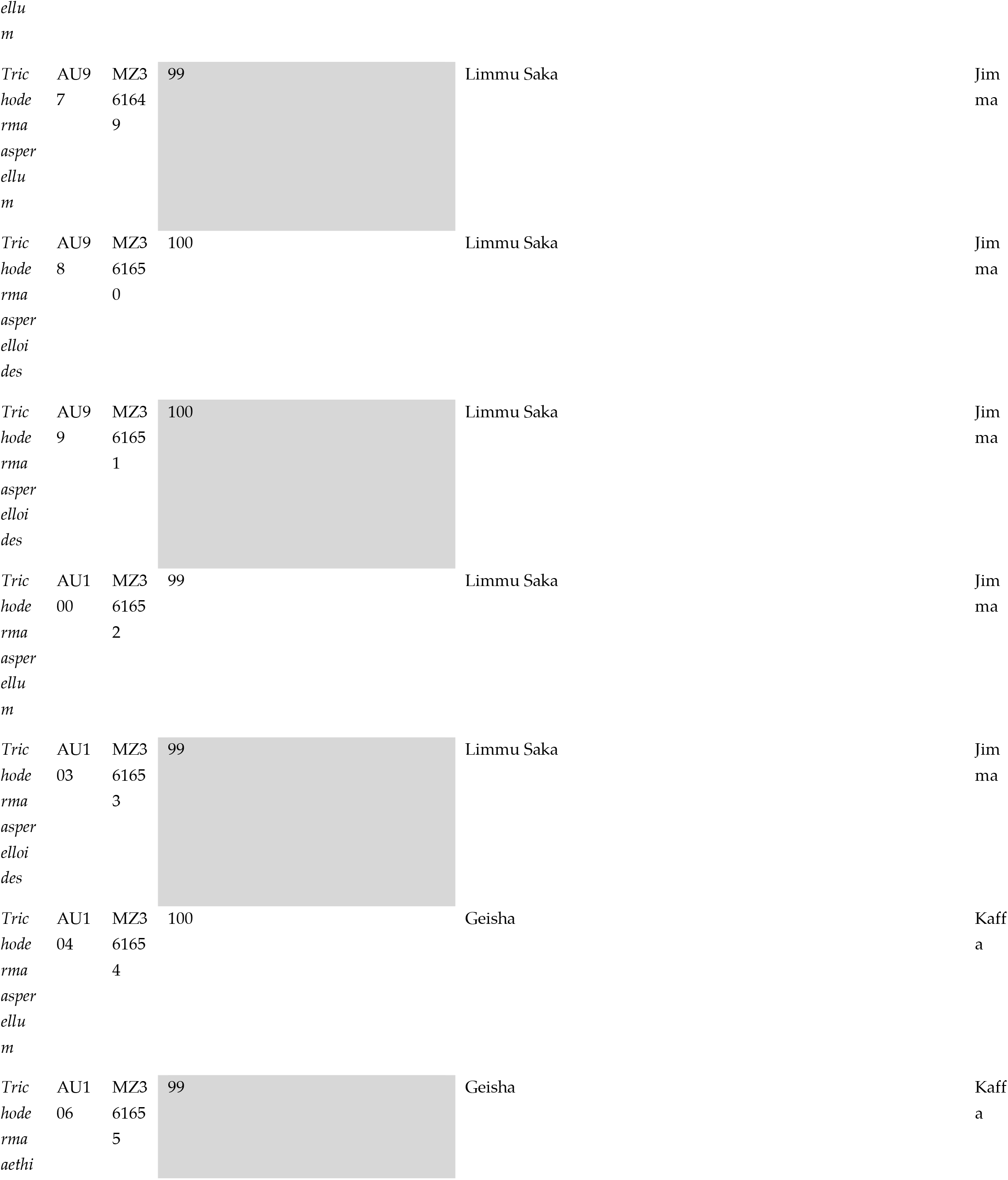
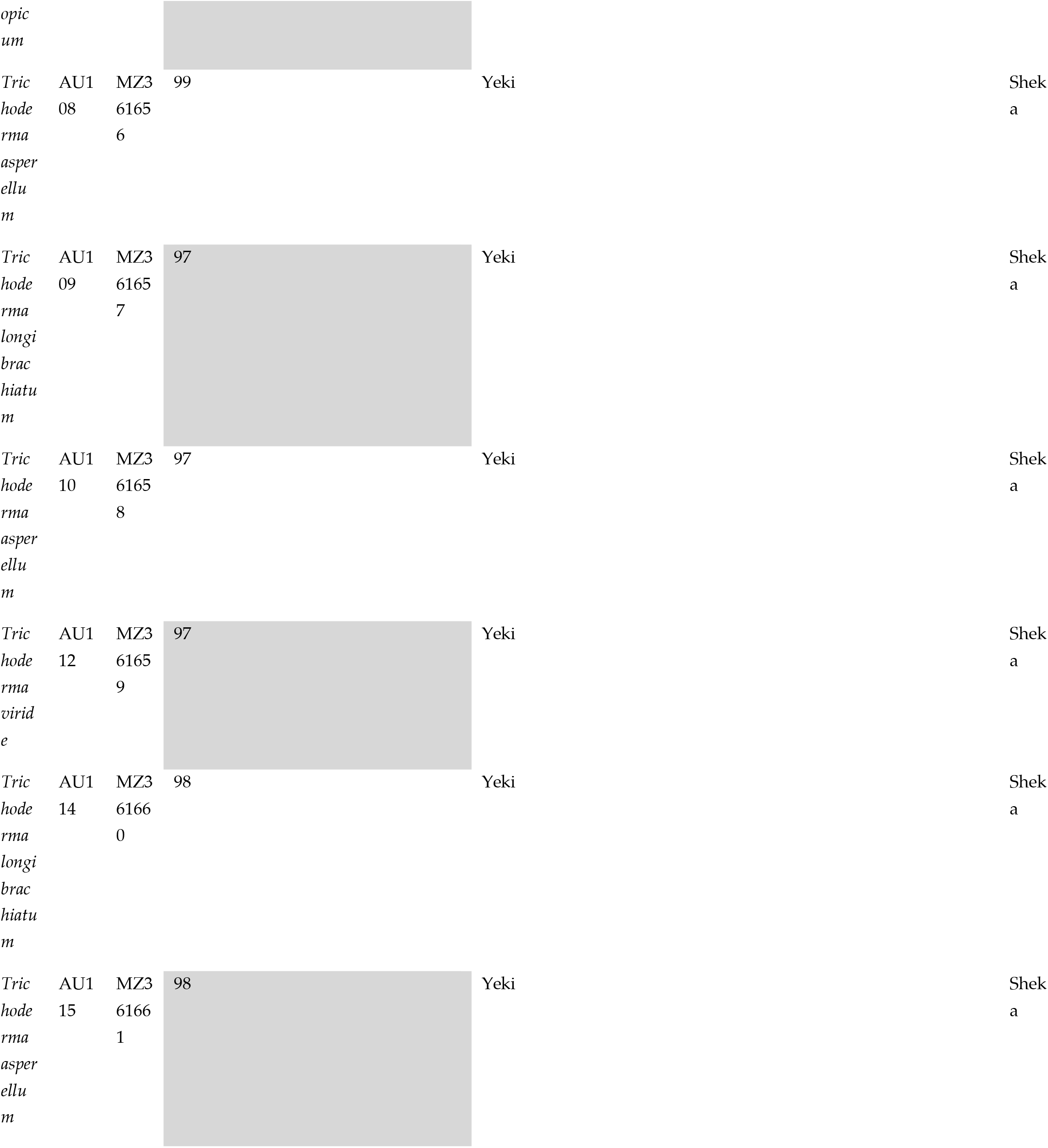
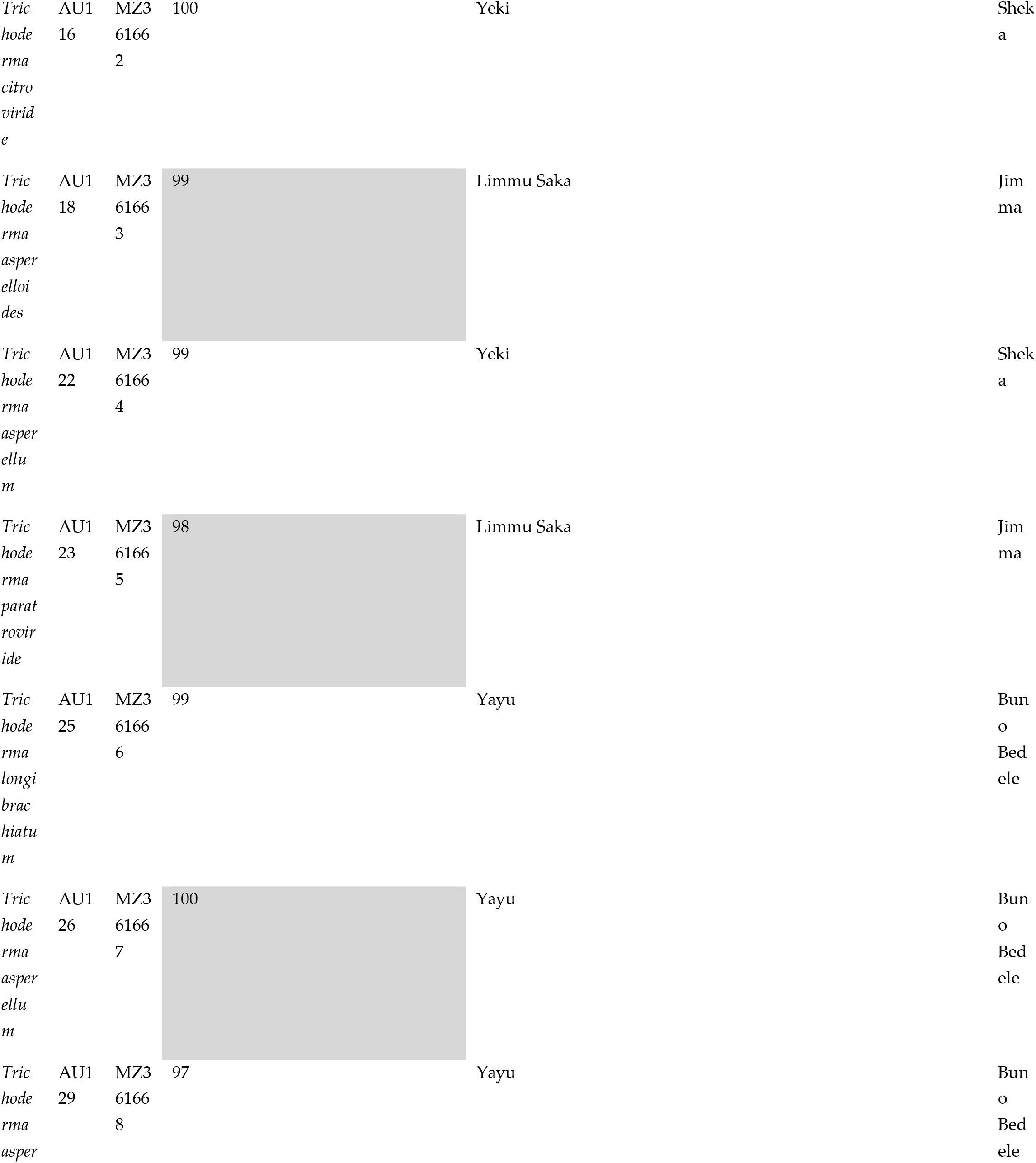
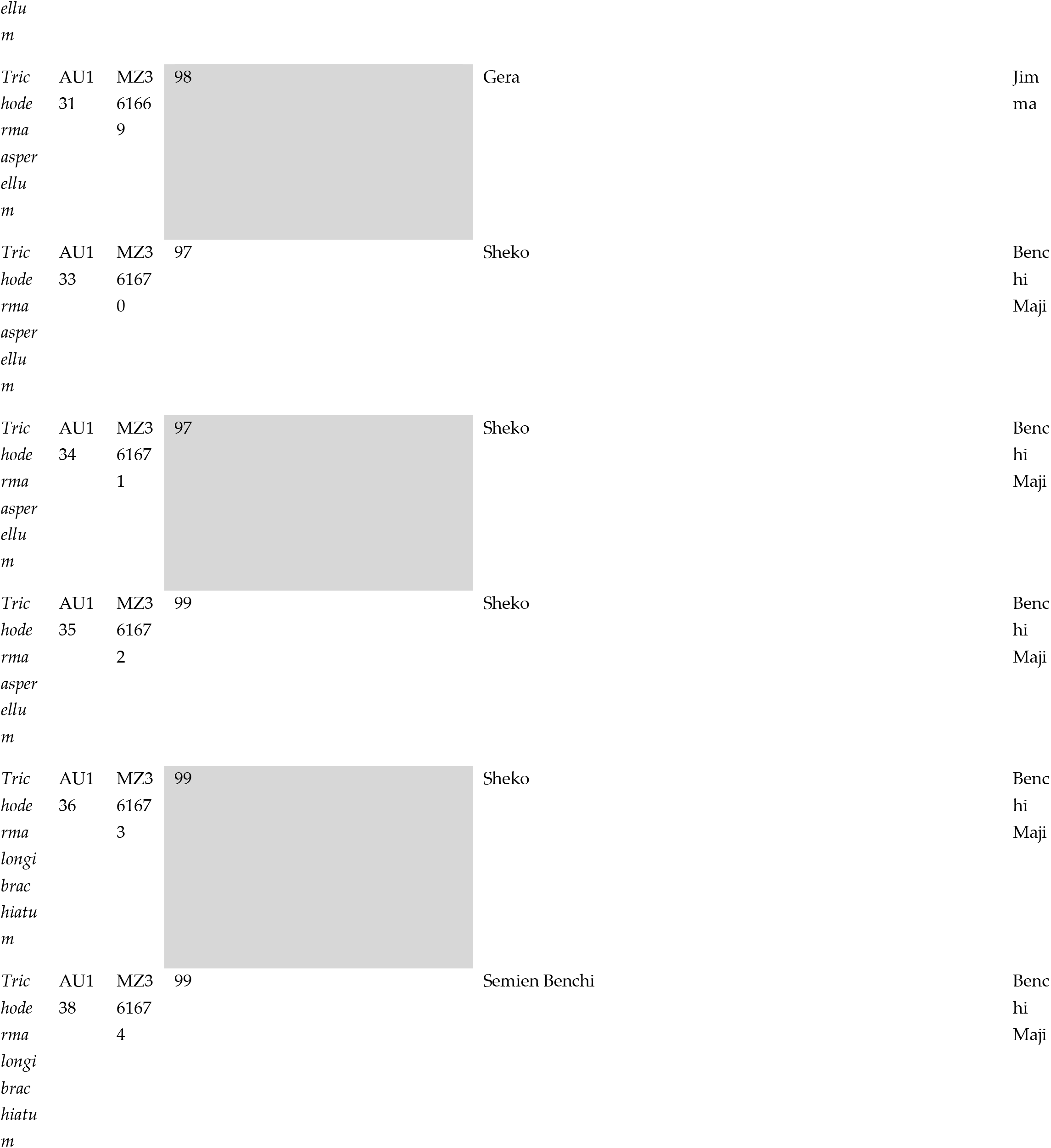
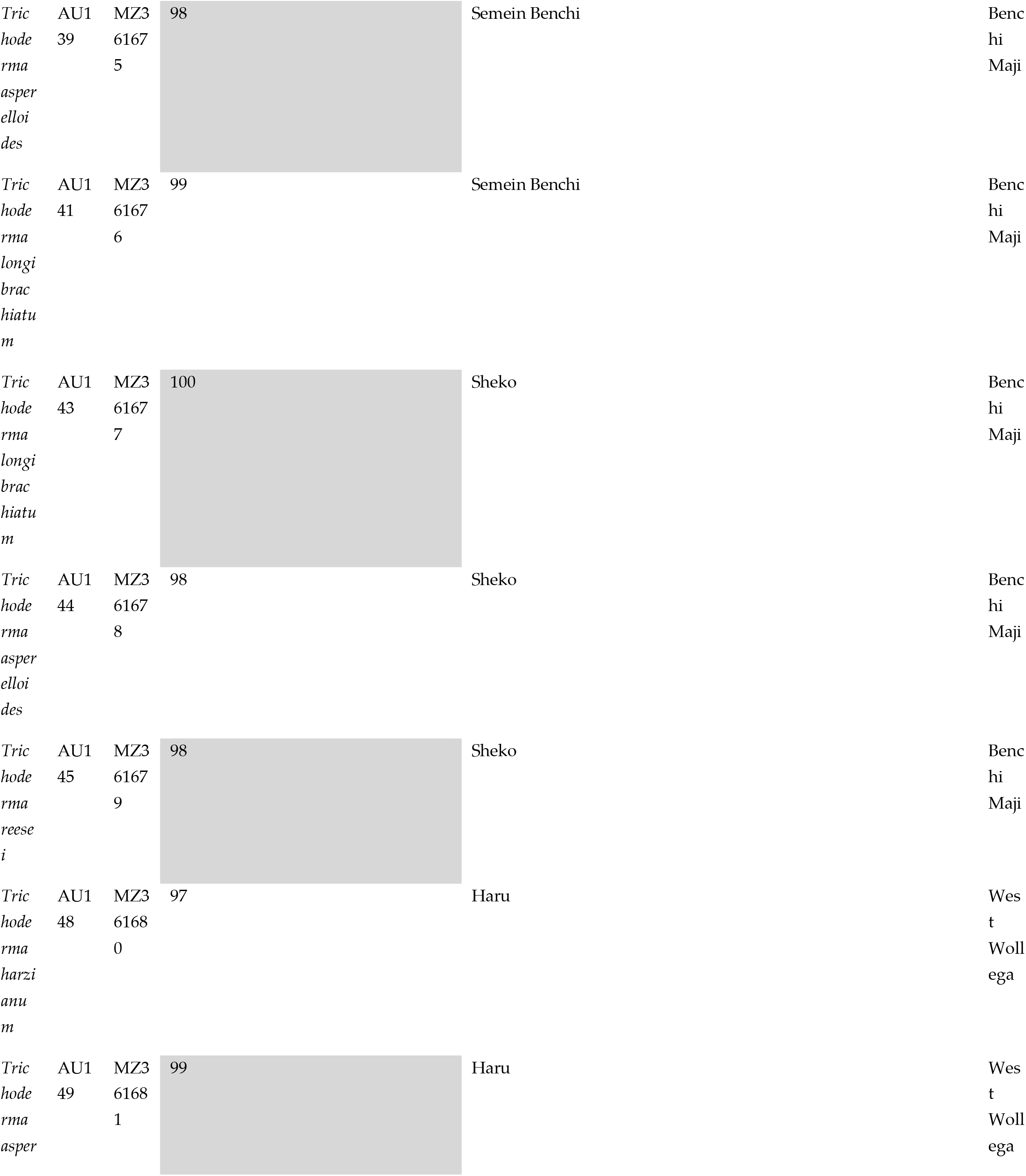
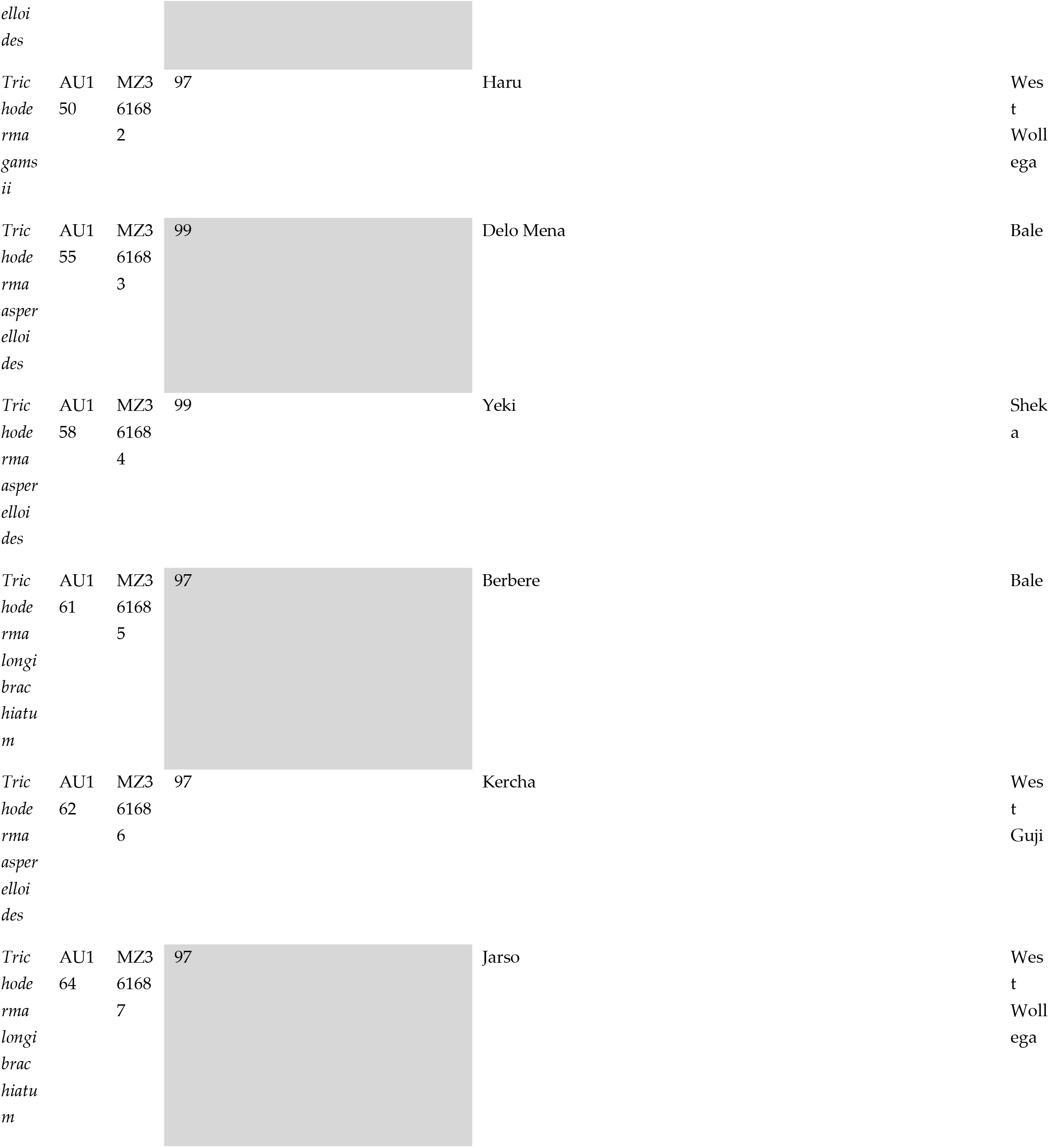
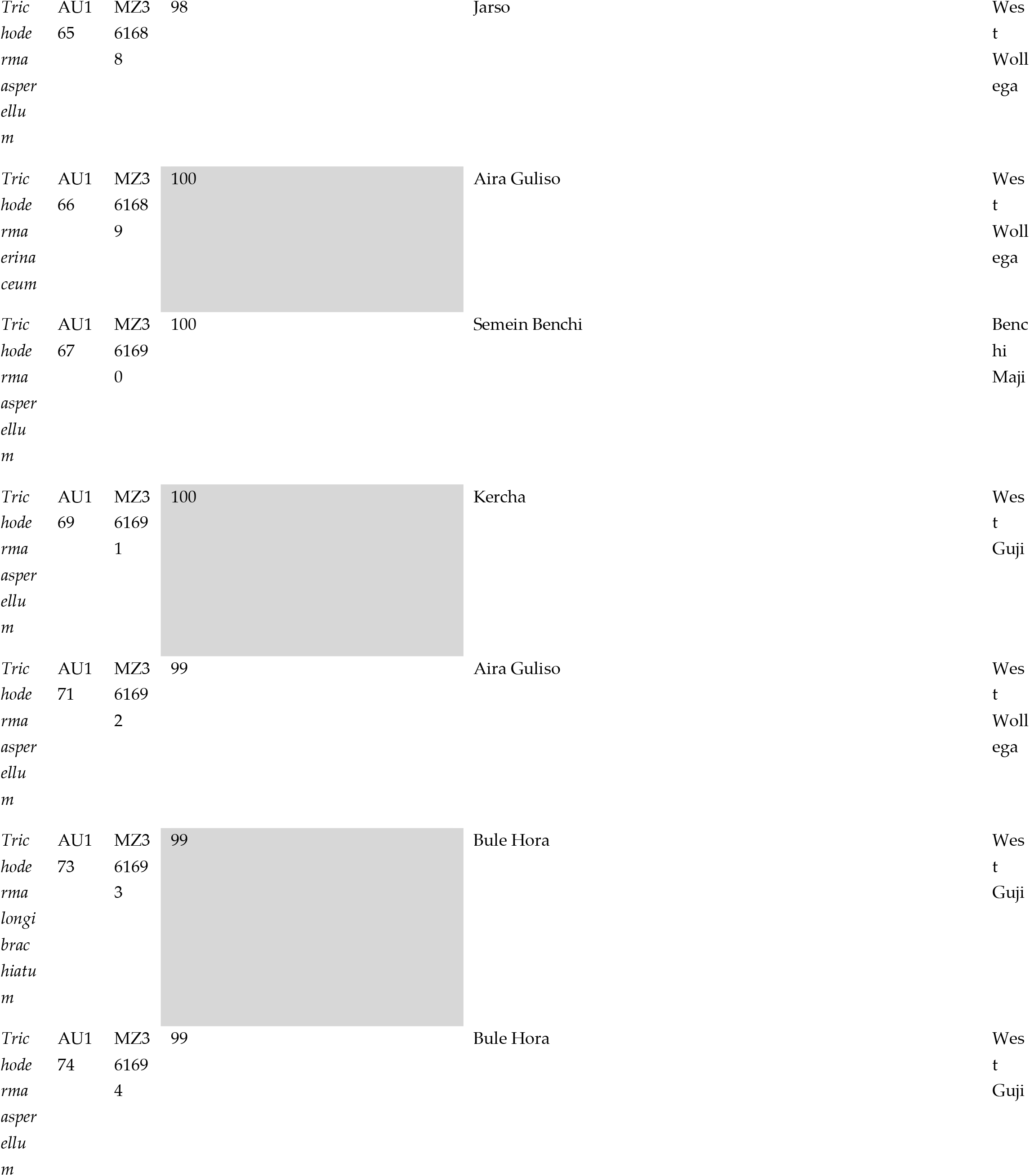
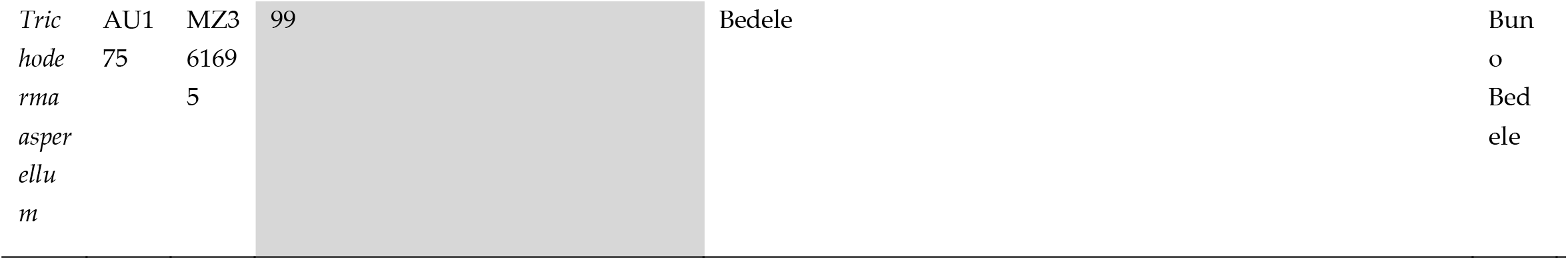
Identification, origin, NCBI Genbank accession numbers, and isolation details of *Trichoderma* species from coffee rhizospheric soil of Ethiopia.

### 2.5 Diversity Analysis of Trichoderma species

The degree of dominance index (Y) was used to quantitatively categorize the habitat preference of *Trichoderma* isolates in the coffee rhizosphere. The dominance values were computed using the following equation:

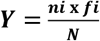

Here, ‘N’ is the total number of *Trichoderma* isolates, ‘n*i*’ is the number of the genus (species) *i*, and ‘*fi*’ is the frequency with which genus (species) *i* appears in the samples. The species *i* is dominant when Y > 0.02 [54]. Species richness (the total number of species), abundance (the sum of the number of isolates of each species), and diversity were evaluated using the Simpson biodiversity index (D) [55], Shannon’s biodiversity index (H) [56], Pielou species evenness index (E) [57], and Margalef’s abundance index (J) [58]. These ecological indices were used to quantitatively describe the diversity and habitat preference of *Trichoderma* species in different coffee ecosystems and major coffee growing zones of Ethiopia.

*Trichoderma* species diversity, defined as the product of the evenness and the number of species, was evaluated using the Shannon biodiversity index (H) [56,59]. Simpson’s diversity index was calculated to assess the dominance of individual species [55,60]. This index shows the probability that two species selected randomly from a given ecosystem will belong to different species categories. Margalef’s abundance index was used to evaluate the species richness while the Pielou index was used to determine the evenness of the *Trichoderma* population. The biological diversity indices were calculated using the following equations:

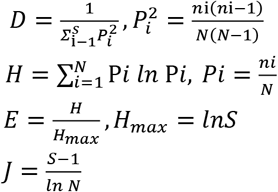

Here, ‘S’ is the total number of *Trichoderma* species, ‘N’ is the sum of all *Trichoderma* species isolates, ‘Pi’ is the relative quantity of *Trichoderma* species ‘i’, and ‘ni’ is the number of isolates of *Trichoderma* species ‘i’.

### 2.6 In vitro Bioassay

In the present study, a total of 175 *Trichoderma* isolates were tested against *F. xylarioides* according to the method of Dennis and Webster [61]. Briefly, mycelial disks (5 mm in diameter) from seven days old growing edges of *Trichoderma* and *F. xylarioides* were put on opposite sides of a PDA Petri dish (3 cm away from each other). Control plates were also prepared without a *Trichoderma* disk. The culture plates were incubated at 25°C with a 12 h photoperiod for 7 days. Following the methodology of [62], the percentage of colonization (%C) of each *Trichoderma* isolate was computed using the formula:

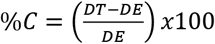

Here, DT is the distance between colonies after mycelial growth stabilizes and DE is the initial distance between the two mycelial discs.

In brief, *Trichoderma* species (1 × 107 spores/ml) were inoculated into 1 liter of PDB at pH 7.2 and cultured for 21 days at 28 °C. After the incubation, the liquid culture was subjected to ethyl acetate extraction and the crude extract was concentrated using a rotary evaporator. Finally, the concentrated extracts were dissolved in methanol for further partial purification using Sephadex LH-20. A total of 25 fractions were collected from the chromatographic column and subjected to agar diffusion assay against *F. xylarioides* on King B medium.

### 2.7 Statistical Data Analysis

Experimental results were analyzed using one-way analysis of variance (ANOVA) with SPSS, version 25. All statistical analyses of ecological indices used to evaluate the biodiversity of *Trichoderma* species were performed using Microsoft Excel 2019 and R software. The significance of differences between the mean results for treatments was evaluated using the Highest Significant Difference (HSD) based on the Tukey test with a significance threshold of *p* ≤ 0.05.

## 3. RESULTS

### 3.1 Isolation and Morphological characterization of Trichoderma Isolates

*Trichoderma* isolates were collected from the coffee rhizosphere conducted in southern, western, and southwestern parts of Ethiopia. A total of 175 *Trichoderma* isolates were obtained from 184 rhizospheric soil samples collected from 28 districts distributed across different agroclimatic zones with soil pH ranging from 4.8 to 8.2. They were morphologically characterized by culturing on PDA plates to capture a full-scale *Trichoderma* diversity and distribution profile. Macroscopic morphological analysis revealed colonies with fast mycelial growth, concentric halos, and floccose or compact surfaces on the culture medium (Figure 2). They were found to form colonies with white mycelia, becoming green when forming conidia and conidiophores. The mycelium, initially of a white color, acquired green or yellow shades, or remained white, due to the abundant production of conidia and secondary metabolites. Concentric rings on culture media were observed for some isolates. Morphological variants including phialides, conidial arrangements, and conidial structures were also observed among the *Trichoderma* isolates. Microscopic analysis revealed plentiful sporulation of conidia originating from verticillate conidiophores. The conidia of most *Trichoderma* isolates were ellipsoidal, globose, subglobose, apex broadly rounded and base more narrowly rounded (Figure 2). However, morphological characteristics were insufficient to distinguish between different *Trichoderma* isolates. Therefore, molecular identification was needed to differentiate the complex and overlapping *Trichoderma* isolates.

**Figure 2.**
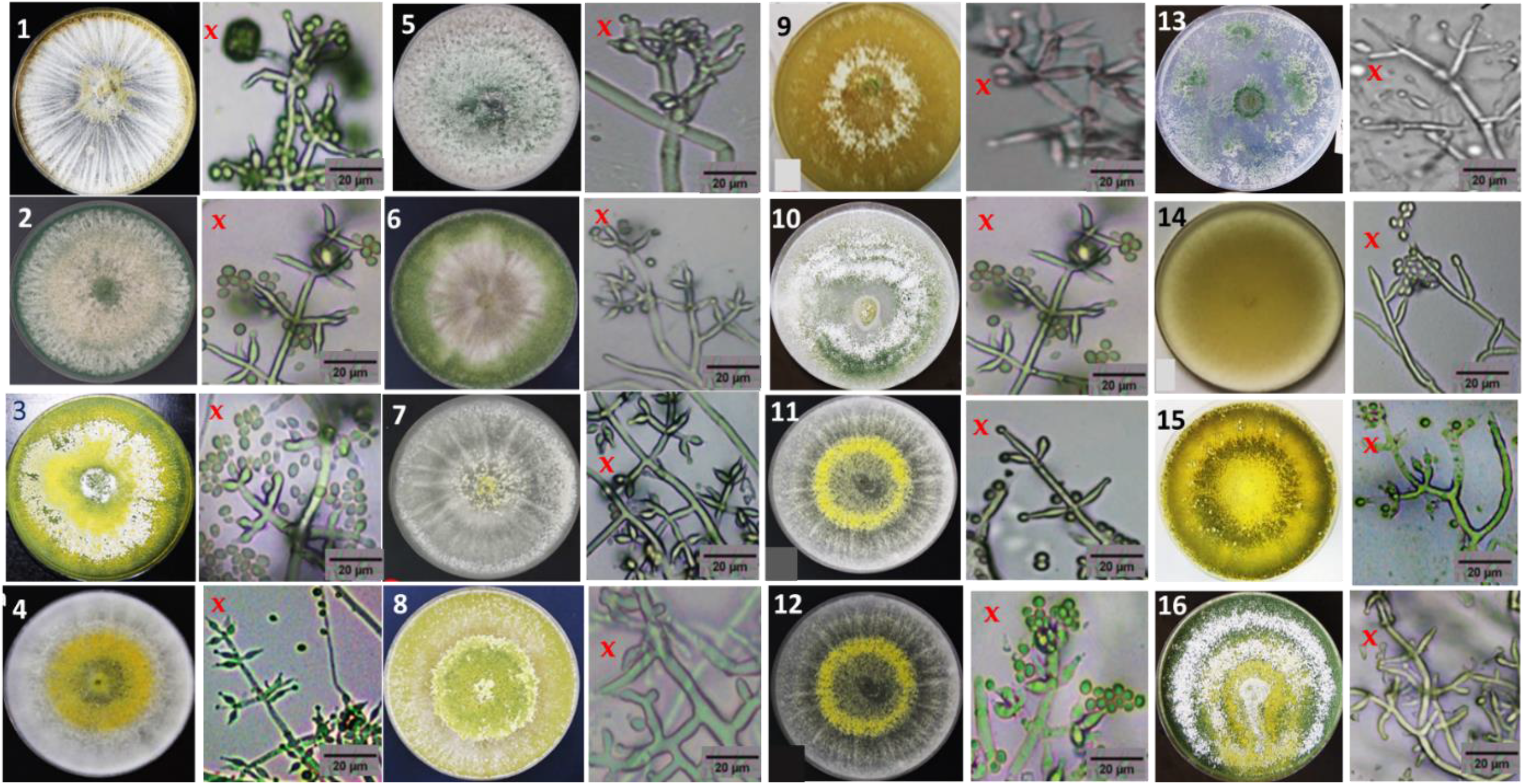
Morphological characteristics of *Trichoderma* species colony grown on PDA: *T. asperellum (*1), *T. asperelloides* (2), *T. longibrachiatum* (3), *T. harzianum* (4), *T. aethiopicum* (5), *T. citrinoviride* (6), *T. hamatum* (7), *T. reesei* (8), *T. vi ride* (9), *T. bissettii* (10), *T. brevicompactum* (11), *T. erinaceum* (12), *T. gamsii* (13), *T. koningiopsis* (14), *T. orientale* (15) and *T. paratroviride* (16); x = structure of conidiophores. Conidiophores were observed with 400× magnification.

### 3.2 Molecular Identification of Trichoderma Isolates

In total, 164 isolates of *Trichoderma* were identified at the species level based on their *TEF1-α* sequences and by morphological analysis. The isolates were assigned to 16 putative species of *Trichoderma*, namely *T. asperellum* (64 isolates), *T. asperelloides* (32), *T. longibrachiatum* (20), *T. harzianum* (8), *T. aethiopicum* (6), *T. hamatum* (6), *T. viride* (4), *T. reesei* (4), *T. koningiopsis* (3), *T. brevicompactum* (3), *T. citrinoviride* (3), *T. gamsii* (3), *T. erinaceum* (2), *T. orientale* (2), *T. bissettii* (3), and *T. paratroviride* (1) (Figure 3 and Supplementary Table S1). These results represent the first observations of the following nine *Trichoderma* species in Ethiopia: *T. asperellum*, *T. bissettii*, *T. brevicompactum, T. citrinoviride*, *T. erinaceum*, *T. orientale, T. paratroviride, T. reesei* and *T. viride.* In addition, 11 undescribed and different isolates could not be matched to any other sequence in Genbank, demonstrating the considerable unresolved biodiversity of *Trichoderma* in the coffee ecosystem.

**Figure 3.**
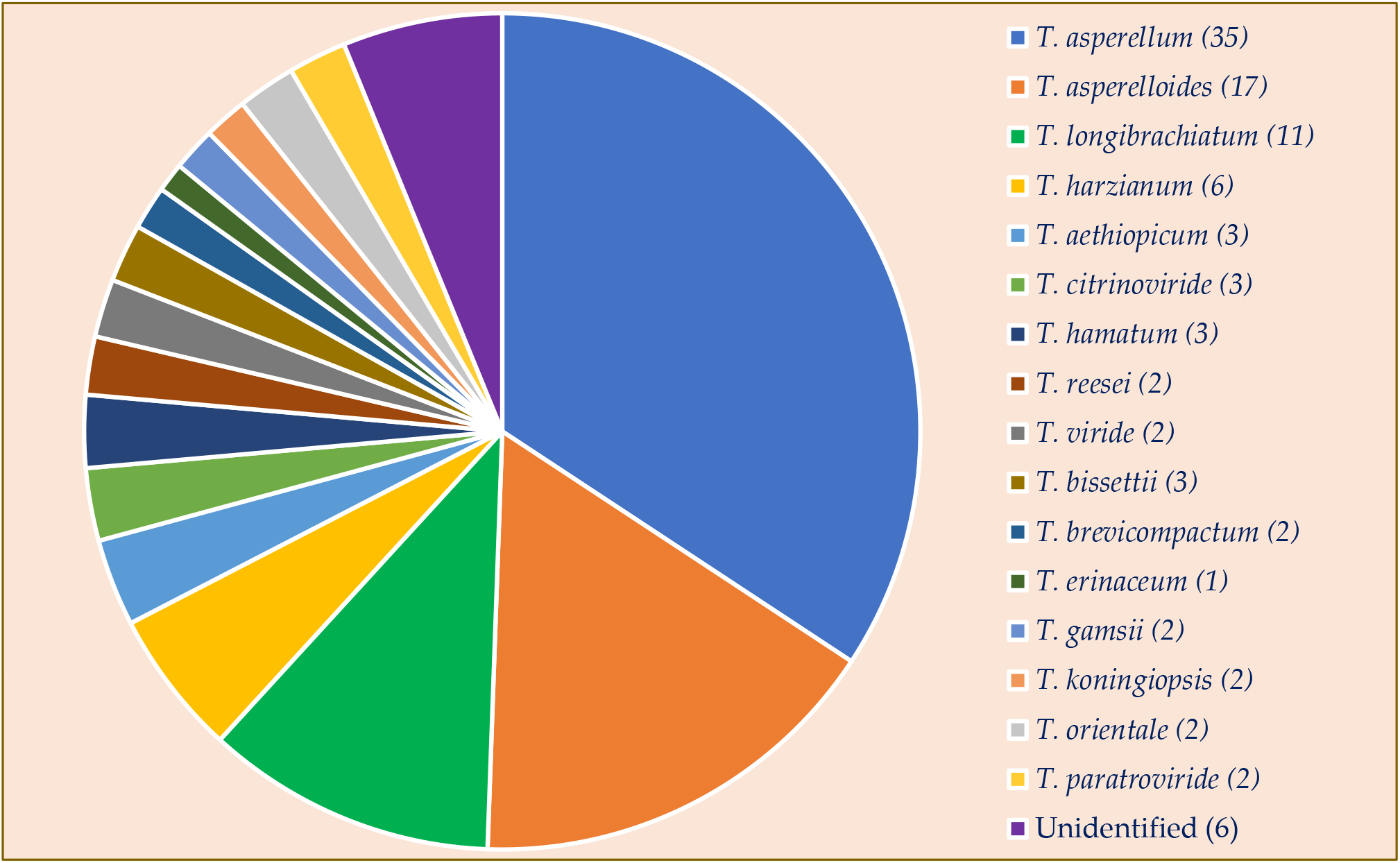
*Trichoderma* species isolated and identified from coffee rhizospheric soil samples: the numbers in parenthesis was the percentage of each *Trichoderma* species.

### 3.3 Phylogenetic Analysis

The *TEF1-α* phylogenetic analysis indicated that the 164 *Trichoderma* isolates were grouped into 16 highly supported monophyletic groups on the phylogeny. The *Tef1-α* phylogenetic analysis and the resulting maximum likelihood tree achieved good resolution for most of the analyzed isolates and effectively discriminated between members of the detected clades. Five basic clades were categorized following the identification manual for *Trichoderma*, namely *Brevicompactum, Longibrachiatum*, *Hamatum*, *Harzianum* and *Viride* (Figure 4). One hundred and two isolates were categorized into three known species belonging to the clades *Hamatum*: *T. asperellum*, *T. asperelloides*, and *T. Hamatum*, while 15 isolates were identified as *T. orientale, T. koningiopsis, T. viride, T. erinaceum, T. paratroviride*, and *T. gamsii* in the clade *Viride.* In addition, 36 isolates were identified as *T. longibrachiatum, T. aethiopicum, T. citroviride, T. bissettii*, and *T. reesei* in the clade *Longibrachiatum*. Eight isolates were identified as *T. harzianum* in the clade *Harzianum*, and 3 isolates grouped as *T. brevicompactum* belonging to the clade *Brevicompactum* (Figure 4).

**Figure 4.** Phylogenetic tree constructed from Maximum Likelihood analysis of *TEF1-α* genes of *Trichoderma*. The *TEF1-α* nucleotide sequences were aligned with similar sequences from taxa of *Trichoderma* species available in the GenBank. The bootstrap scores are based on 1,000 iterations. The scale bar represents 50 substitutions per nucleotide position. Sequences from this study are designated with isolate ID: AU.

### 3.4 Biodiversity and Distribution of Trichoderma Isolates

#### 3.4.1 Diversity analysis of *Trichoderma* species

The dominance value (Y) was 0.048 (> 0.02), indicating that the genus *Trichoderma* was dominant in coffee rhizosphere soil. *T. asperellum*, *T. asperelloides* and *T. longibrachiatum* were classified as the principal species, with dominance (Y) values of 0.062, 0.056, and 0.034, respectively. The analysed data were used to compute Simpson’s biodiversity index (D), Shannon’s biodiversity index (H), evenness (E), and the abundance index (J) for each coffee ecosystem and coffee growing zone, as shown in Supplementary Table S1. The highest species diversity and evenness (H = 1.97, E = 0.79, D = 0.81) were recorded in the forest and semi-forest coffee ecosystems of Kaffa, Jimma, and Bale. The Shannon and Simpson diversity indices estimated for the garden coffee ecosystems of the West Guji and Bunno Bedele zones showed that they had lower species diversity (H = 1.57, D = 0.7). The calculated species abundance values were E = 2.71 for forest coffee, E = 2.64 for semi-forest coffee, and E = 2.14 for garden coffee. The average diversity values for *Trichoderma* species originating from the coffee ecosystem were H = 1.77, D = 0.7, E = 0.75 and J = 2.4 (Table 2). Simpson’s index and the evenness index were close to 1 except in the West Guji zone, indicating a very high diversity of *Trichoderma* species in major coffee growing areas of Ethiopia. The numbers of species and isolates, and the dominant species of *Trichoderma* varied geographically (Table S2). These results revealed that the forest, semi-forest, and garden ecosystems had a high diversity of *Trichoderma* species. The rhizosphere of *C*. *arabica* in Ethiopia thus hosts a large and highly diverse population of *Trichoderma* species.

**Table 2.** Univariate diversity indices analysis of *Trichoderma* isolates in different coffee ecosystems and major coffee growing zones of Ethiopia.

| Ecological indices | Coffee Ecosystem |  |  |  | Major coffee growing zones |  |  |  |  |  |  |  |  |  |
| --- | --- | --- | --- | --- | --- | --- | --- | --- | --- | --- | --- | --- | --- | --- |
|  | Native Forest | Semi forest | Garden coffee | Average | Jimma | Kaffa | Bench Maji | Sheka | Bunno Bedale | West Wollega | West Guji | Gedio | Sidama | Bale |
| Simpson index (D) | 0.81 | 0.81 | 0.7 | 0.76 | 0.91 | 0.94 | 0.83 | 0.82 | 0.7 | 0.64 | 0.53 | 0.9 | 0.8 | 0.87 |
| Shannon's index (H) | 1.97 | 1.96 | 1.57 | 1.77 | 1.97 | 1.82 | 1.7 | 1.83 | 0.75 | 0.94 | 1.29 | 1.33 | 1.54 | 1.89 |
| Pielou evenness Index (E) | 0.79 | 0.79 | 0.71 | 0.75 | 0.76 | 0.95 | 0.87 | 0.83 | 0.86 | 0.72 | 0.68 | 0.96 | 0.86 | 0.97 |
| Abundance index (J) | 2.71 | 2.64 | 2.14 | 2.4 | 2.73 | 2.58 | 1.97 | 2.49 | 1.24 | 1.95 | 1.17 | 1.86 | 2.09 | 2.6 |

#### 3.4.2 Distribution of *Trichoderma* species in different coffee growing zones

Distribution and habitat preference analysis showed that *Trichoderma* species were widely dispersed throughout different coffee production systems. The proportion and composition of *Trichoderma* species varied among the sampled coffee growing districts and zones. The proportion of *Trichoderma* species obtained from Jimma zone was the highest (27%), followed by Sheka zone (16%) and Bench Maji zone (13%); the lowest proportion was obtained in the Bunno Bedele zone (3%) (Figure 5). Species richness was highest in the Jimma zone (25 soil samples), from which 11 *Trichoderma* species were identified, followed by the Sheka zone (9 species, 18 soil samples), whereas Bunno Bedele had only 3 *Trichoderma* species. Among the identified isolates, *T. asperellum* (39.6%) and *T. asperelloides* (28%) were the most abundant species, being found in all major coffee growing zones and districts of Ethiopia (Figure 5). Conversely, *T. paratroviride* was noted only in soil samples collected from Jimma zone. The number of *Trichoderma* species declined on going from the southwest to the south. The 11 known species identified in the Jimma zone were *T. asperellum*, *T. asperelloides*, *T. longibrachiatum*, *T. harzianum*, *T. aethiopicum, T. citrinoviride*, *T. viride*, *T. reesei, T. koningiopsis*, *T. erinaceum* and *T. paratroviride*. On the other hand, *Trichoderma* species obtained from the Sheka zone were *T. asperellum*, *T. asperelloides*, *T. longibrachiatum*, *T. viride*, *T. hamatum, T. brevicompactum, T. koningiopsis*, *T. citrinoviride,* and *T. bissettii*. *T. asperellum* and *T. asperelloides* were found in all major coffee growing areas and were the most widely dispersed species. Another widely distributed species was *T. longibrachiatum*, which was scattered in all zones except Kaffa. However, some species were unique to one zone; for instance, *T. paratroviride* was isolated only from Jimma zone (Figure 5).

**Figure 5.**
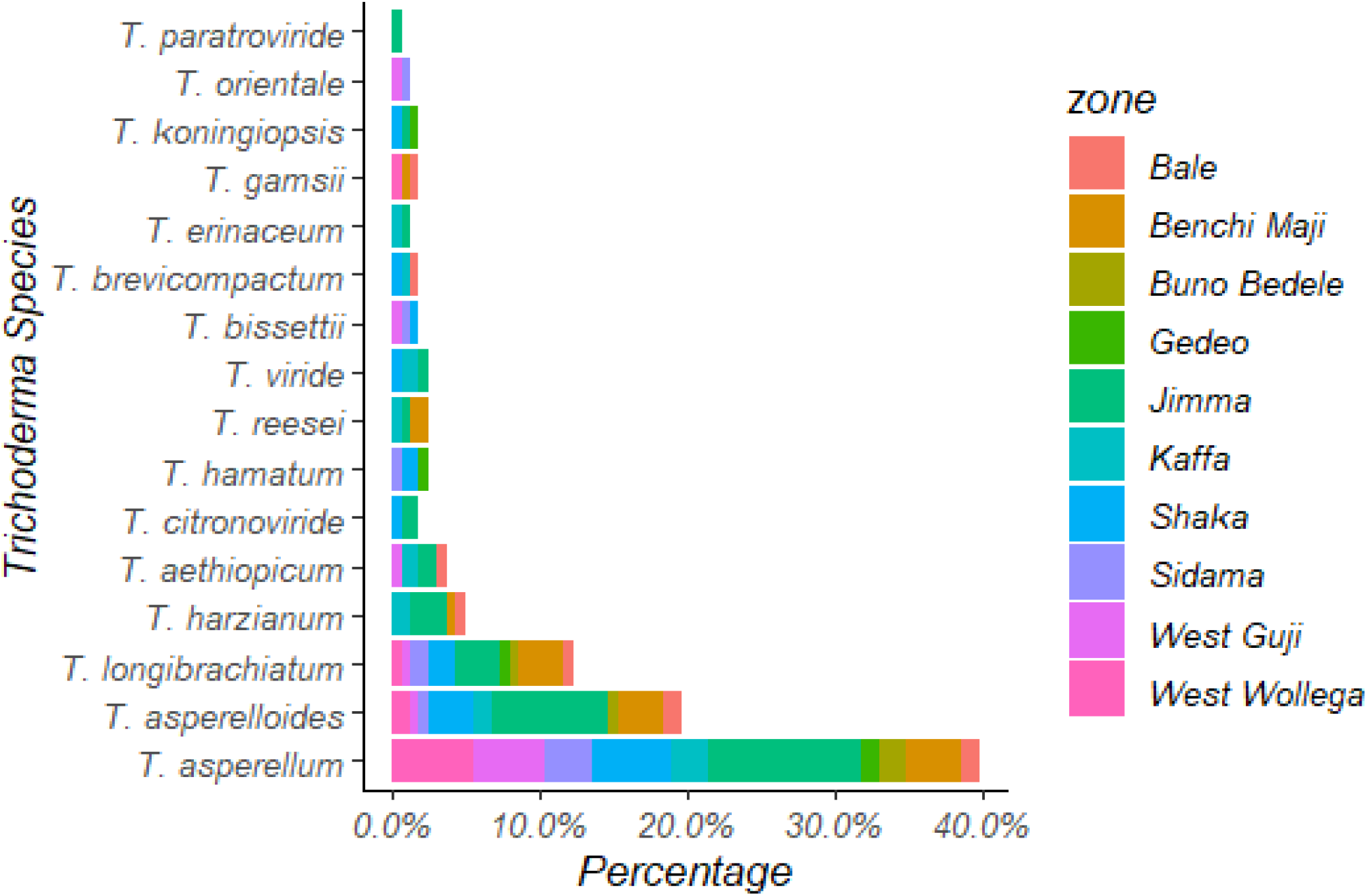
Distribution of *Trichoderma* species in major coffee growing zones of Ethiopia.

#### 3.4.3 Distribution of *Trichoderma* species in a Coffee Ecosystem

There were slight differences in the communities of *Trichoderma* species observed in the coffee rhizosphere soils of the different coffee ecosystems. Their high biodiversity was apparent in the distribution of *Trichoderma* species (Table S1). In total, 72 soil samples were collected from the native forest ecosystem, yielding 68 isolates representing 12 species of *Trichoderma*. Fifty-nine soil samples were collected from disturbed semi-forests, yielding 62 isolates representing 13 species. Fewer samples were collected from garden coffee ecosystems (53 soil samples), yielding only 9 different *Trichoderma* species. The isolation frequency of *Trichoderma* in the native forest ecosystem was 39%, which was substantially higher than that for garden coffee ecosystems (29%; Figure 6). Except for species represented by single isolates, all species were found in multiple areas, showing that they may be regularly distributed within the coffee rhizosphere. However, there were some notable exceptions: *T. erinaceum* and *T. brevicompactum* were mostly isolated from the forest rhizosphere, *T. paratroviride* and *T. citrinoviride* were only found in semi forest zones, and *T. orientale* was only observed in the garden coffee ecosystem (Figure 6).

**Figure 6.**
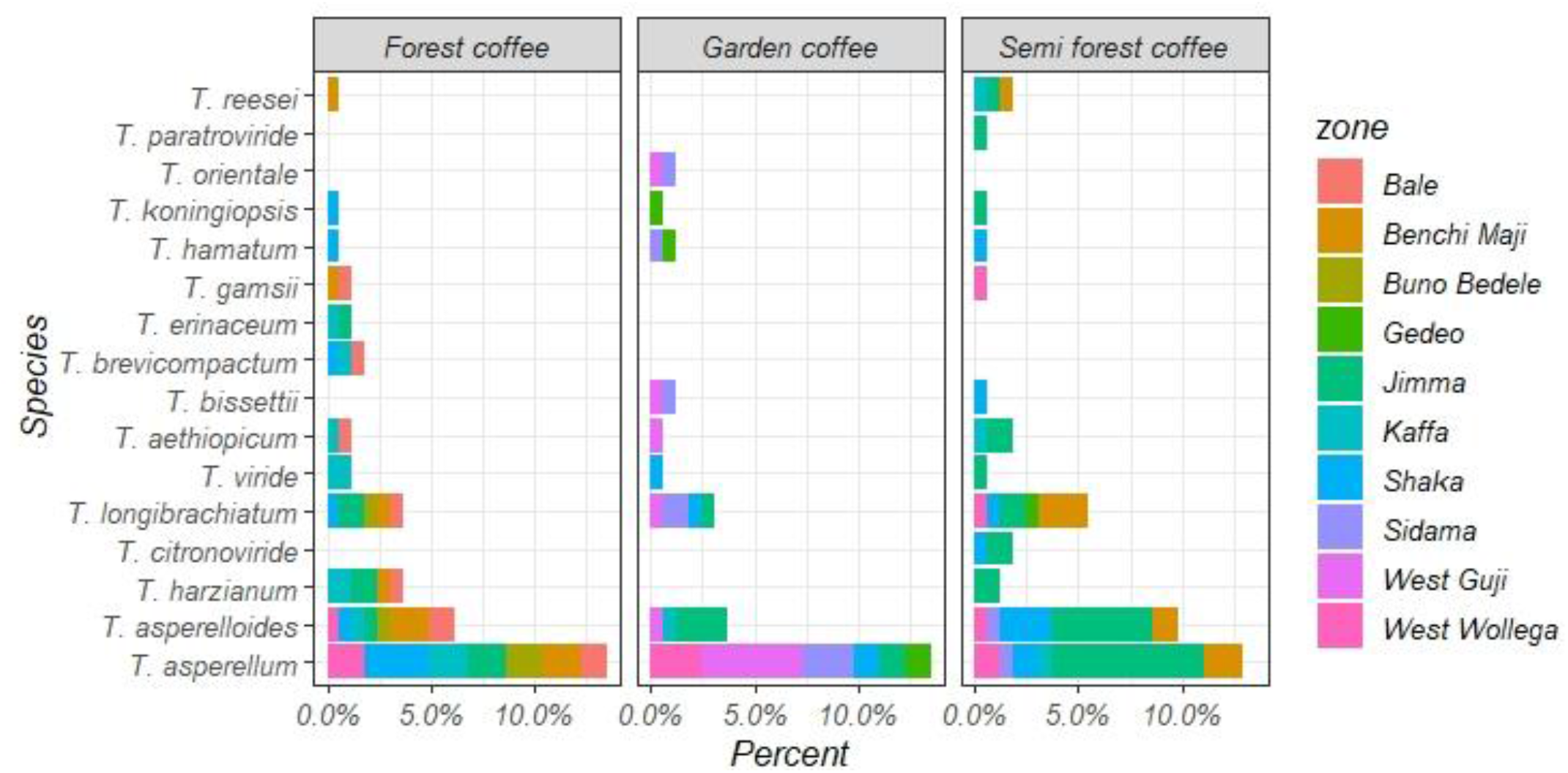
Distribution of *Trichoderma* species in different coffee production ecosystems.

### 3.5 Screening of Biocontrol Trichoderma

All isolates were capable of significantly inhibiting the mycelial growth of *F. xylarioides*. Twelve isolates exhibited the highest defined level of *in vitro* antagonistic activity. ANOVA analysis revealed statistically significant (*p* ≤ 0.05) differences in the mycelial growth inhibition profiles of the *Trichoderma* isolates against *F. xylarioides*, with inhibition percentages ranging from 44.5% to 84.8% (Table 3). Among them, *T. asperellum* AU71, *T. longibrachiatum* AU158 and *T. asperellum* AU131 were the most effective, causing 79.3%, 82.4%, and 84.8% inhibition, respectively ((Table 3 and Figure 7 A-C)). The mean inhibitory effect of these isolates against *F. xylarioides* was such that the pathogen’s growth was suppressed almost completely whereas it grew rapidly on control plates lacking *Trichoderma* isolates ((Figure 7 (1D)). The inhibition of *F. xylarioides* radial growth in the dual culture confronting assay was attributed to inhibitory secondary metabolites released by one or both organisms as well as competition, mycoparasitism, and production of cell wall degrading enzymes. The potential *Trichoderma* species exhibited an average growth rate of 0.45 mm/h in dual culture bioassays.

**Table 3.** *In vitro* evaluation of *Trichoderma* isolates against *F. xylarioides* by dual confrontation culture technique and agar diffusion assay.

| *Trichoderma*<br>species | Mycelia Inhibition over control (%) |  | Scale of antagonistic activity |
| --- | --- | --- | --- |
|  | Dual Culture | Agar diffusion assay |  |
| *T. hamatum* AU23 | 71<sup>c</sup> ± 1.07 | 67.32<sup>ab</sup> ± 4.06 | +++ |
| *T. longibrachiatum* AU32 | 75.2<sup>b</sup> ± 0.7 | 69.82<sup>ab</sup> ± 4.20 | +++ |
| *T. asperellum* AU53 | 72.6<sup>b</sup> ± 0.3 | 66.3<sup>c</sup> ± 2.3 | +++ |
| *T. koningiopsis* AU70 | 62.59<sup>d</sup> ± .9 | 70.71<sup>ab</sup> ± 4.82 | +++ |
| *T. asperellum* AU71 | 79.3<sup>a</sup> ± 3.03 | 83.5<sup>a</sup> ± 4.83 | ++++ |
| *T. asperellum* AU97 | 74.8<sup>b</sup> ± 1.0 | 76.42<sup>b</sup> ± 3.68 | ++++ |
| *T. harzianum* AU105 | 68.7<sup>c</sup> ± 1 | 75.82<sup>b</sup> ± 4.81 | +++ |
| *T. aethiopicum* AU106 | 59.3<sup>d</sup> ± 5.1 | 68.50<sup>c</sup> ± 5.12 | ++ |
| *T. longibrachiatum* AU121 | 79.2<sup>b</sup> ± 0.9 | 63.4<sup>c</sup> ± 3.4 | +++ |
| *T. asperellum* AU131 | 84.8<sup>a</sup> ± 0.9 | 86.7<sup>a</sup> ± 1.6 | ++++ |
| *T. longibrachiatum* AU158 | 82.4<sup>a</sup> ± 0.5 | 88.2<sup>a</sup> ± 3.5 | ++++ |
| *T. asperellum* AU171 | 77.7<sup>ab</sup> ± 0.3 | 66.4<sup>c</sup> ± 2.5 | +++ |
| Mean ± standard deviation | 73.96 ± 1.3 | 74.25 ± 3.74 | - |

**Figure 7.**
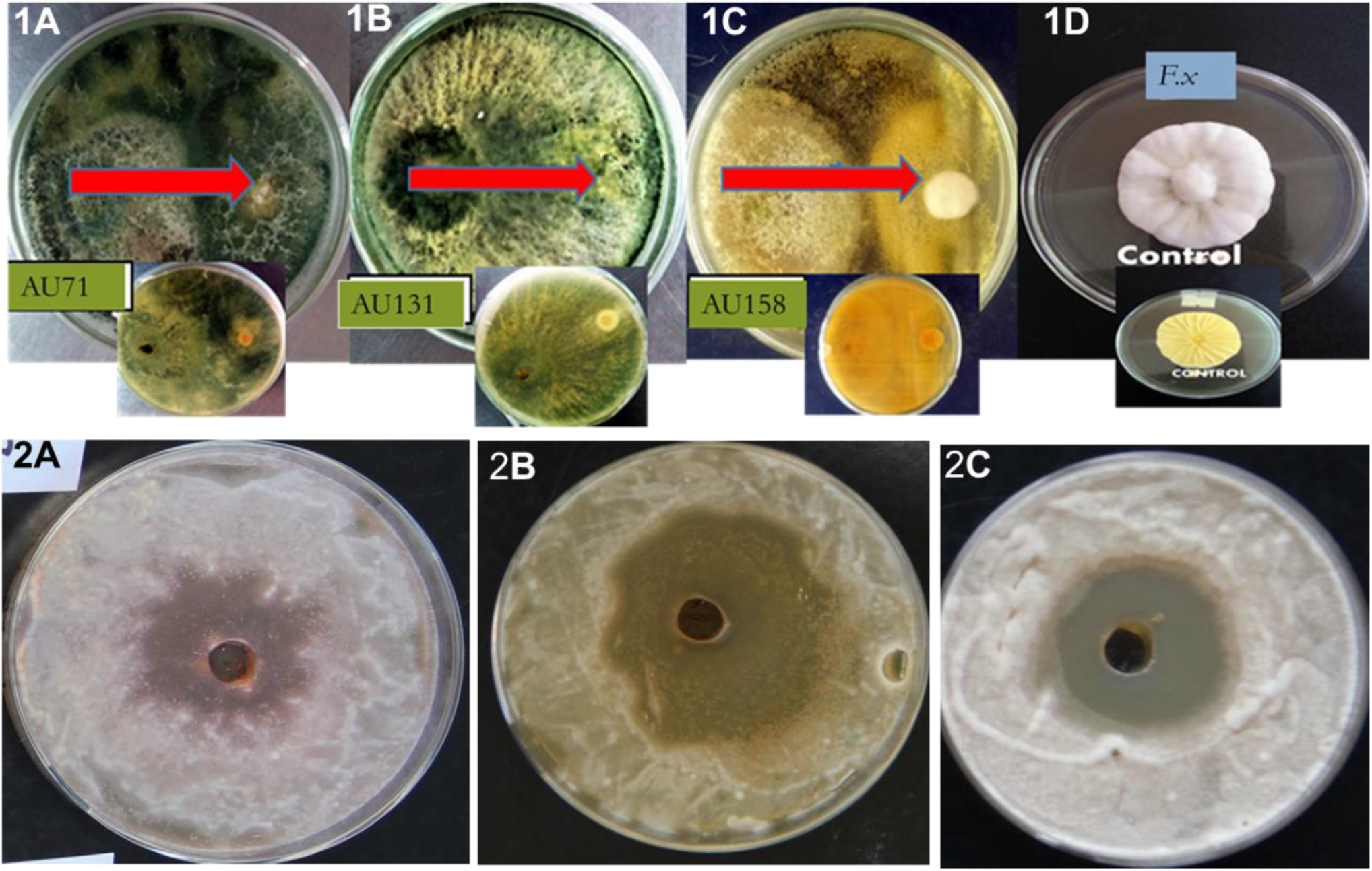
Antagonistic effects of *Trichoderma* species against *F. xylarioides:* (1A-C) dual culture bioassay, (2A-C) agar diffusion bioassay. *T. asperellum* AU71 (A), *T. asperellum* AU131 (B*), T. longibrachiatum* AU158 (C) and *F. xylarioides (*1D) alone as a control. Red arrows indicate the growth of the test pathogen.

Based on the *in vitro* bioassay results, three potent isolates (*T. asperellum* AU71, *T. asperellum* AU131 and *T. longibrachiatum* AU158) were subjected to secondary metabolite extraction. The agar well diffusion method was used to quantify the antifungal activities of crude metabolites extracted from these species ((Table 3 and Figure 7 (2A-C)). All crude metabolites from these microorganisms inhibited the mycelial growth of *F. xylarioides* at the point of application around the agar wells; inhibition percentages of 83.5%, 86.7%, and 88.2% were observed for the extracts of *T. asperellum* AU71, *T. asperellum* AU131, and *T. longibrachiatum* AU158, respectively, ((Figure 7 (2ABC)) (*p* ≤ 0.05).

## 4. Discussion

A total of 164 isolates belonging to five clades were obtained from coffee rhizosphere soil samples. *Trichoderma* species were primarily identified based on morphological characteristics including green coloration interleaved with a white mycelium, which is consistent with the morphological features reported previously for this fungus [31,63]. The identification keys of Samuels*, et al.* [63] and Rifai [31] state that *T. longibrachiatum* holds subglobous to ovoid conidia and lageniform phialides. Additionally, [35] describes the presence of yellowish-green pigment on the backside of plates of *T. longibrachiatum*; which was also observed in this work. However, phenotypic characters are varied and depend partly on culture conditions [64] and secondary metabolite production [65]. This plasticity of characteristics means that analyses based solely on phenotypic traits cannot provide conclusive taxonomic identification of *Trichoderma* species [66,67].

Phylogenetic grouping revealed that the *Trichoderma* isolates recovered in this study formed a reliable maximum likelihood tree with acceptable taxonomic assumptions [68,69]. Modern methodologies for *Trichoderma* identification and classification into phylogenetic clades are based on analyses of sequence data [41,67,69]. Five clades were identified in this study, namely; *Hamatum*, *Harzianum*, *Longibrachiatum*, *Brevicompactum* and *Viride* (Figure 4). The *Hamatum* clade contains economically important species such as *T. asperellum* and *T. asperelloides*, which are used in agriculture as biological control agents [70,71]. The *Longibrachiatum* clade has high optimal and maximum growth temperatures and yellow reverse pigmentation due to the production of secondary metabolites such as pyrone. *Trichoderma longibrachiatum* has been used to produce various antimicrobial substances with important agricultural, health, and environmental benefits [19].

The diversity of *Trichoderma* species in Africa in general [43,72], and in Ethiopia in particular [24,73] is somewhat understudied when compared to other parts of the world. Nine *Trichoderma* species were identified for the first time in Ethiopia in this work. It is notable that these species were previously described in America [74], Asia [46,54,75] and in European Mediterranean countries [42,76]; their presence in coffee rhizosphere soils in Ethiopia can be attributed to the diverse ecological substrata and climate conditions of the country’s coffee growing areas and reflects the high *Trichoderma* biodiversity present in coffee ecosystems. The only previous study comparable to this one in terms of sampling size and studied area was conducted in the neotropical forests of South America, mainly in Colombia [77]. In that study, a high diversity of *Trichoderma* (29 species among 183 isolates) was detected, with a high proportion of putative new species among the isolates (11 species, corresponding to 6% of the sample). The main difference between their findings and ours is that we investigated a well-defined microecological niche, namely the rhizosphere of *C*. *arabica.*

The biodiversity of *Trichoderma* species is difficult to evaluate comparatively due to range of indices suggested for this purpose [78]. In the present study, several widely used diversity indices were tested using a range of simple and multifaceted statistical analyses to evaluate whether some were better suggested for certain analyses than others. The Shannon index values calculated for native forest and semi-forest ecosystem samples were almost twice those obtained for soils in Sardinia; H = 1.97 *versus* 1.59, respectively, even though the number of samples investigated in the latter case was almost three times that collected in this work. However, the Shannon indices of the Sardinian ecosystems and the garden coffee zones were quite similar (H = 1.59 *versus* 1.57), possibly reflecting the extensive disturbance of both ecosystems by human activities [79]. These results showed that *Trichoderma* diversity and habitat preference can be used as a natural indicator of rhizosphere soil health. Forest and semi-forest coffee regions had richly varied *Trichoderma* populations with relatively high diversity and very similar biodiversity indices and evenness values.

The number of *Trichoderma* species detected in this work was almost twice that reported in earlier studies on biodiversity in Ethiopia [24] and other countries including Poland [80], Central Europe [76], and China’s Northern Xinjiang region [75]. In addition, significant differences were observed between the *Trichoderma* populations of different coffee growing zones; this variation may reflect differences in the zones’ ecological environments. The populations of *Trichoderma* species in the forest and semi-forest coffee ecosystems of southwestern Ethiopia were diverse and their composition varied between ecosystems. Jimma zone had 11 *Trichoderma* species and the largest number of *Trichoderma* isolates (48), followed by the Sheka (9 species, 27 isolates), Benchi Maji (7 species, 22 isolates), and Bunno Bedele (3 species, 6 isolates) zones (Table 1). Our results suggest that forest and semi-forest ecosystems are particularly favorable for the survival and colonization of *Trichoderma*, indicating that this genus has a clear environmental preference, in keeping with previous reports [54,75,76].

*T. asperellum* (39%) was found to be the most widely distributed and abundant fungal species in this work (Figure 3). The occurrence of *Trichoderma* species is modulated by several factors, including metabolic variety, reproductive ability, substrate availability, and the competitive abilities of *Trichoderma* isolates in nature [76,81,82]. *Trichoderma* isolates were obtained from different coffee ecosystems, with *T. asperellum, T. asperelloides* and *T. longibrachiatum* being the most widely distributed species. *T. asperellum* is the most dominant and cosmopolitan species like *T. harzianum* [83], whereas *T. asperelloides* and *T. longibrachiatum* were found mostly in forest ecosystems of South America and Asia [77,84]. Conversely, previous studies have found *T*. *harzianum*, *T*. *hamatum*, *T*. *spirale,* and *T*. *asperelloides* to be the most widely distributed species of this genus in coffee ecosystem in Ethiopia [24]. Except for species that were only found as single isolates, all species were obtained in multiple districts, suggesting that they are quite evenly distributed within the coffee rhizosphere. *T. erinaceum* and *T. brevicompactum* were only isolated from the native forest; *T. paratroviride* and *T. citrinoviride* were only obtained from semi-forest areas, and *T. orientale* was only isolated from garden coffee ecosystem samples. Studies conducted by Hoyos-Carvajal and Bissett [77] indicated the dominant *Trichoderma* species in the neotropics are *T. asperellum*, followed by *T. harzianum*. Our results confirmed the predominance of *T. asperellum*, followed by *T. asperelloides*. Conversely, Belayneh*, et al.* [24] reported that *T. hamatum* was the most dominant species in the rhizosphere of coffee plants. The large number and wide distribution of *Trichoderma* species identified within Ethiopia’s coffee ecosystem demonstrate the presence of significant genetic diversity, suggesting that further study of these species may offer opportunities to improve the sustainable management of coffee cultivation and discover effective bio-control agents for managing CWD.

This work represents the first investigation of the biodiversity of *Trichoderma* species in the rhizospheres of Ethiopia’s coffee ecosystem and their suitability as biological control agents (BCA) against CWD (*F. xylarioides*). The results presented herein mainly concern the taxonomy of the *Trichoderma* isolates with some observations on their ecology, and will support the selection of candidate biocontrol agents for the management of CWD in Ethiopia. This work is part of a larger project seeking to control CWD using a classical biological control strategy involving sourcing and releasing potential BCA from the center of origin of *coffea arabica* to minimize the incidence and severity of the disease. Such approaches using fungal natural enemies have been used successfully to control various soil-borne plant pathogens [9,24,73,85,86]. Our results indicate that there is a lot of *Trichoderma* species that are substantially antagonistic to *F. xylarioides* and which could be exploited for the biocontrol of CWD in this way. In the previous study, we formulated a biofungicide from *T*. *asperellum* AU131 and *T. longibrachiatum* AU158 under solid state fermentation (SSF) to control of CWD [87].

All *Trichoderma* strains isolated in this work effectively inhibited the mycelial growth of *F. xylarioides* colonies. However, there were notable differences between strains in terms of the magnitude of their antagonism and reduction of the growth rate of *F. xylarioides* in paired culture experiments. For example, Filizola*, et al.* [88] state that some isolates of certain species suppress the growth of phytopathogens via hyper-parasitism, whereas others achieve growth suppression via mechanisms such as antibiosis or competition. It has also been reported that *Trichoderma* species grow faster than competing phytopathogens because they use food sources more efficiently. Another important mechanism involves the secretion of metabolites and hydrolytic enzymes that reduce or hinder the growth of plant pathogens in the area; this mechanism has been suggested to contribute to the success of *Trichoderma* species against phytopathogenic fungi [89]. The potential of *T. asperellum* and *T. longibrachiatum* as effective biocontrol agent of fungi and bacterial strains of both on annual and perennial crops were clearly stated by many research reports [90,91], For instance, *T. asperellum* exhibits strong control effects on *F. graminearum*, *F. oxysporum and Verticillium* wilt of olive [92,93]. On the other hand, *T. longibrachiatum* is also used as potential biocontrol agents most effective against *P. grisea*, *F. verticillioides*, *H. oryzae*, *F. moniliforme*, and *A. alternate* with inhibition percentages of 98.9, 96.4, 95.1, 93.6, and 93.0%, respectively [94]. We should point out here the fact that the three *T. asperellum* strains assessed in this work were isolated from coffee rhizosphere in production fields from southwestern Ethiopia. This aspect should be considered a valuable asset for biocontrol applications, as native isolates are better adapted to their local climate conditions and pathogenic targets than foreign isolates.

Various secondary metabolites produced by *Trichoderma* species including harzianolide, peptaibols, pyrones and other secondary metabolites were mentioned to have antimicrobial potential and to act as plant growth promoters (Mulatu, unpublished data). In addition to achieving higher growth rates than *F. xylarioides* in competition experiments, the *Trichoderma* strains isolated in this work achieved growth rates significantly exceeding the value of 0.33 mm/h reported by [95]. Moreover, field and greenhouse experiments using Geisha coffee varieties of *C*. *arabica* (the most susceptible to CWD) gave similar results (Afrasa Mulatu., unpublished data). The results obtained indicate that understanding the genetic variation within the genus *Trichoderma* is essential for selecting novel indigenous *Trichoderma* species that can be used as biocontrol agents against CWD. In addition, our findings display the distribution and diversity profile of *Trichoderma* species and provide insights into their potential usefulness as microbial fungicides to safeguard coffee cultivation across different agroclimatic zones in Ethiopia.

## 5. Conclusions

A total of 175 isolates of *Trichoderma* were identified at the species level based on *TEF1-α* sequence analysis, yielding 16 putative species*. T. asperellum*, *T. asperelloides*, and *T. longibrachiatum* were classified as the abundant species and the average diversity values for *Trichoderma* species originating from coffee ecosystems were H = 1.77, D = 0.7, E = 0.75, and J = 2.4. The results obtained suggest that *T. asperellum* and *T. longibrachiatum* are promising suppressors of *F. xylarioides* growth and promoters of plant growth, suggesting that they could be valuable biocontrol agents for the management of CWD. Additionally, our results demonstrated the existence of a guild of *Trichoderma* species that are potentially antagonistic to *F. xylarioides*, which could be exploited to develop more effective biological control of CWD.

## Supporting information

Table S1. Trichoderma species isolated and identified from major coffee growing zones of Ethiopia and Table S2. Identification, origin, and isolation

## Author Contributions

Conceptualization and designing the experiment, A.M., R.R.V., and T.A.; methodology, A.M., T.A. and N.M.; DNA extraction, T.A and A.M, PCR optimization and sequencing, R.R.V, S.K and Q.L; software, A.M.; Data validation, T.A. and R.R.V.; formal analysis, A.M.; investigation, A.M. and T.A.; resources, T.A., N.M. and R.R.V.; data curation, A.M. and T.A.; writing-original draft preparation, A.M.; writing-review and editing, T.A., N.M. and R.R.V.; supervision and project administration, T.A., N.M. and R.R.V.; funding acquisition, T.A., and R.R.V. All authors have read and agreed to the published version of the manuscript.

## Institutional Review Board Statement

Not applicable.

## Data Availability Statement

Not applicable.

## Acknowledgments

We would like to thank EBTi for funding this research project under the theme “Bio-fungicide production for CWD control in Ethiopia” and Department of Microbial, Cellular and Molecular Biology of Addis Ababa University (AAU), Ethiopia, for the laboratory facilities. Special thanks goes to the Department of Plant Breeding of the Swedish University of Agricultural Sciences (SLU), Sweden, for valuable support to the project. RV was supported by the Swedish Research Council for Environment, Agricultural Sciences and Spatial Planning (FORMAS) (grant number 2019-01316) and the Swedish Research Council (grant number 2019-04270).

