## Supplementary material for "Biodiversity of the Genus *Trichoderma* in the Rhizosphere of Coffee (*Coffea arabica*) Plants in Ethiopia and their Potential Use in Biocontrol of Coffee Wilt Disease": Table S1. Trichoderma species isolated and identified from major coffee growing zones of Ethiopia and Table S2. Identification, origin, and isolation

| *Trichoderma* species | Jimma | Kaffa | Benchi Maji | Shaka | Buno Bedele | West Wollega | West Guji | Gedio | Sidama | Bale | Total isolates |
|---|---|---|---|---|---|---|---|---|---|---|---|
| Soil samples | 25 | 20 | 23 | 18 | 16 | 14 | 17 | 16 | 21 | 14 | 184 |
| *T. asperellum* | 16 | 4 | 6 | 9 | 3 | 9 | 8 | 2 | 5 | 2 | 64 |
| *T. asperelloides* | 13 | 2 | 5 | 5 | 1 | 2 | 1 | | 1 | 2 | 32 |
| *T. longibrachiatum* | 4 | 0 | 5 | 4 | 1 | 1 | 1 | 1 | 2 | 1 | 20 |
| *T. harzianum* | 4 | 2 | 1 | 0 | 0 | 0 | 0 | 0 | 0 | 1 | 8 |
| *T. aethiopicum* | 2 | 2 | 0 | 0 | 0 | 0 | 1 | 0 | 0 | 1 | 6 |
| *T. citrinoviride* | 2 | 0 | 0 | 1 | 0 | 0 | 0 | 0 | 0 | 0 | 3 |
| *T. hamatum* | 1 | 0 | 1 | 2 | 0 | 0 | 0 | 1 | 1 | 0 | 6 |
| *T. reesei* | 1 | 1 | 2 | 0 | 0 | 0 | 0 | 0 | 0 | 0 | 4 |
| *T. viride* | 1 | 2 | 0 | 1 | 0 | 0 | 0 | 0 | 0 | 0 | 4 |
| *T. bissettii* | 0 | 0 | 0 | 1 | 0 | 0 | 1 | 0 | 1 | 0 | 3 |
| *T. brevicompactum* | 0 | 1 | 0 | 1 | 0 | 0 | 0 | 0 | 0 | 1 | 3 |
| *T. erinaceum* | 1 | 1 | 0 | 0 | 0 | 0 | 0 | 0 | 0 | 0 | 2 |
| *T. gamsii* | 0 | 0 | 1 | 0 | 0 | 1 | 0 | 0 | 0 | 1 | 3 |
| *T. koningiopsis* | 1 | 0 | 0 | 1 | 0 | 0 | 0 | 1 | 0 | 0 | 3 |
| *T. orientale* | 0 | 0 | 0 | 0 | 0 | 0 | 1 | 0 | 1 | 0 | 2 |
| *T. paratroviride* | 1 | 0 | 0 | 0 | 0 | 0 | 0 | 0 | 0 | 0 | 1 |
| Non- identified | 1 | 2 | 1 | 2 | 1 | 1 | 0 | 3 | 0 | 0 | 0 |
| ***Total*** | 48 | 17 | 22 | 27 | 6 | 14 | 13 | 8 | 11 | 9 | 175 |

Table S2. Identification, origin, and isolation details of *Trichoderma* isolates

| **Isolate ID** | **Taxa** | **District (Woreda)** | **Zone** | **Coffee Ecosystem** |
|---|---|---|---|---|
| AU1 | *T. asperellum* | Gera | Jimma | Semi forest |
| AU2 | *T. hamatum* | Gera | Jimma | Semi forest |
| AU3 | *T. asperellum* | Gera | Jimma | Semi forest |
| AU4 | *T. citrinoviride* | Gera | Jimma | Semi forest |
| AU5 | *T. asperelloides* | Gera | Jimma | Semi forest |
| AU6 | *T. reesei* | Melko | Jimma | Semi forest |
| AU7 | *T. koningiopsis* | Gera | Jimma | Semi forest |
| AU8 | *T. asperellum* | Gera | Jimma | Semi forest |
| AU9 | *T. longibrachiatum* | Yeki | Jimma | Semi forest |
| AU10 | *T. aethiopicum* | Gera | Jimma | Semi forest |
| AU11 | *T. asperelloides* | Gera | Jimma | Semi forest |

| | | | | |
|---|---|---|---|---|
| AU12 | *T. aethiopicum* | Gera | Jimma | Semi forest |
| AU13 | *T. asperellum* | Gera | Jimma | Semi forest |
| AU14 | *T. longibrachiatum* | Yeki | Shaka | Semi forest |
| AU15 | *T. asperellum* | Gera | Jimma | Semi forest |
| AU16 | *T. harzianum* | Gera | Jimma | Semi forest |
| AU17 | *T. asperellum* | Gera | Jimma | Semi forest |
| AU18 | *T. asperellum* | Gera | Jimma | Semi forest |
| AU19 | *T. citrinoviride* | Gera | Jimma | Semi forest |
| AU20 | *T. asperellum* | Limmu Saka | Jimma | Semi forest |
| AU21 | *T. asperellum* | Gera | Jimma | Semi forest |
| AU22 | *T. asperellum* | Yeki | Shaka | Semi forest |
| AU23 | *T. viride* | Gera | Jimma | Semi forest |
| AU24 | *T. orientale* | Odo Shakiso | West Guji | Garden Coffee |
| AU25 | *T. asperellum* | Odo Shakiso | West Guji | Garden Coffee |
| AU26 | *T. asperellum* | Odo Shakiso | West Guji | Garden Coffee |
| AU27 | *T. asperellum* | Odo Shakiso | West Guji | Garden Coffee |
| AU28 | *T. asperelloides* | Gera | Jimma | Garden Coffee |
| AU29 | *T. asperelloides* | Gera | Jimma | Garden Coffee |
| AU30 | *T. hamatum* | Shebedino | Sidama | Garden Coffee |
| AU31 | *T. asperellum* | Shebedino | Sidama | Garden Coffee |
| AU32 | *T. longibrachiatum* | Gera | Jimma | Garden Coffee |
| AU33 | *T. asperellum* | Shebedino | Sidama | Garden Coffee |
| AU34 | *T. asperelloides* | Yeki | Shaka | Forest |
| AU35 | *T. asperellum* | Yeki | Shaka | Forest |
| AU36 | *T. longibrachiatum* | Melko | Jimma | Forest |
| AU37 | *T. erinaceum* | Gera | Jimma | Forest |
| AU38 | *T. asperellum* | Yeki | Shaka | Forest |
| AU39 | *T. asperellum* | Gimbo | Kaffa | Forest |
| AU40 | *T. longibrachiatum* | Gomma | Jimma | Forest |
| AU41 | *T. brevicompactum* | Yeki | Shaka | Forest |
| AU42 | *T. asperellum* | Gomma | Jimma | Forest |
| AU43 | *T. asperellum* | Haru | West Wollega | Forest |
| AU44 | *T. asperellum* | Gomma | Jimma | Forest |
| AU45 | *T. asperelloides* | Gomma | Jimma | Forest |
| AU46 | *T. asperellum* | Gomma | Jimma | Forest |
| AU47 | *T. asperelloides* | Chena | Kaffa | Forest |
| AU48 | *T. koningiopsis* | Yeki | Shaka | Forest |
| AU49 | *T. longibrachiatum* | Andaracha | Shaka | Forest |
| AU50 | *T. asperellum* | Andaracha | Shaka | Forest |
| AU51 | *T. hamatum* | Andaracha | Shaka | Forest |
| AU52 | *T. asperellum* | Andaracha | Shaka | Forest |
| AU53 | *T. asperellum* | Andaracha | Shaka | Forest |
| AU54 | *T. asperellum* | Mena | Jimma | Semi forest |
| AU55 | *T. asperelloides* | Mena | Jimma | Semi forest |
| AU56 | *Non identified* | Gewata | Kaffa | Forest |
| AU57 | *T. asperelloides* | Mena | Jimma | Semi forest |
| AU58 | *T. erinaceum* | Gewata | Kaffa | Forest |
| AU59 | *T. bissettii* | Aleta Wondo | Sidama | Garden Coffee |
| AU60 | *T. longibrachiatum* | Aleta Wondo | Sidama | Garden Coffee |
| AU61 | *T. asperelloides* | Gomma | Kaffa | Garden Coffee |
| AU62 | *T. asperellum* | Jarso | West Wollega | Garden Coffee |

| | | | | |
|---|---|---|---|---|
| AU63 | *T. asperellum* | Jarso | West Wollega | Garden Coffee |
| AU64 | *Non identified* | Yirga cheffe | Gedeo | Garden Coffee |
| AU65 | *Non identified* | Yirga cheffe | Gedeo | Garden Coffee |
| AU66 | *Non identified* | Yirga cheffe | Gedeo | Garden Coffee |
| AU67 | *T. longibrachiatum* | Aleta Wondo | Sidama | Garden Coffee |
| AU68 | *T. asperellum* | Yirga cheffe | Gedeo | Garden Coffee |
| AU69 | *T. asperellum* | Wonago | Gedeo | Garden Coffee |
| AU70 | *T. koningiopsis* | Wonago | Gedeo | Garden Coffee |
| AU71 | *T. asperelloides* | Yirga cheffe | Sidama | Semi forest |
| AU72 | *T. longibrachiatum* | Yirga cheffe | Gedeo | Semi forest |
| AU73 | *T. asperellum* | Gewata | Kaffa | Semi forest |
| AU74 | *T. asperellum* | Aleta Wondo | Sidama | Semi forest |
| AU75 | *T. asperellum* | Dale | Sidama | Garden |
| AU76 | *T. asperellum* | Dale | Sidama | Garden |
| AU77 | *T. orientale* | Dale | Sidama | Garden |
| AU78 | *T. harzianum* | Gimbo | Kaffa | Forest |
| AU79 | *T. viride* | Gimbo | Kaffa | Forest |
| AU80 | *T. asperellum* | Haru | West Wollega | Forest |
| AU81 | *T. asperellum* | Haru | West Wollega | Forest |
| AU82 | *T. asperellum* | Sheko | Benchi Maji | Forest |
| AU83 | *T. asperellum* | Sheko | Benchi Maji | Forest |
| AU84 | *T. harzianum* | Sheko | Benchi Maji | Forest |
| AU85 | *T. asperelloides* | Sheko | Benchi Maji | Forest |
| AU86 | *T. gamsii* | Sheko | Benchi Maji | Forest |
| AU87 | *T. harzianum* | Gera | Jimma | Forest |
| AU88 | *T. harzianum* | Gera | Jimma | Forest |
| AU89 | *T. asperelloides* | Gera | Jimma | Semi forest |
| AU90 | *T. viride* | Chena | Kaffa | Forest |
| AU91 | *T. asperellum* | Chena | Kaffa | Forest |
| AU92 | *T. brevicompactum* | Chena | Kaffa | Forest |
| AU93 | *T. reesei* | Chena | Kaffa | Semi forest |
| AU94 | *T. aethiopicum* | Chena | Kaffa | Semi forest |
| AU95 | *T. asperellum* | Limmu Saka | Jimma | Semi forest |
| AU96 | *T. asperellum* | Limmu Saka | Jimma | Semi forest |
| AU97 | *T. asperellum* | Limmu Saka | Jimma | Garden Coffee |
| AU98 | *T. asperelloides* | Limmu Saka | Jimma | Garden Coffee |
| AU99 | *T. asperelloides* | Limmu Saka | Jimma | Garden Coffee |
| AU100 | *T. asperellum* | Limmu Saka | Jimma | Garden Coffee |
| AU101 | *Non identified* | Limmu Saka | Jimma | Semi forest |
| AU102 | *T. hamatum* | Yeki | Shaka | Semi forest |
| AU103 | *T. asperelloides* | Limmu Saka | Jimma | Semi forest |
| AU104 | *T. asperellum* | Geisha | Kaffa | Forest |
| AU105 | *T. harzianum* | Geisha | Kaffa | Forest |
| AU106 | *T. aethiopicum* | Geisha | Kaffa | Forest |
| AU107 | *Non identified* | Geisha | Kaffa | Forest |
| AU108 | *T. asperelloides* | Yeki | Shaka | Semi forest |
| AU109 | *T. bissettii* | Yeki | Shaka | Semi forest |
| AU110 | *T. asperellum* | Yeki | Shaka | Semi forest |
| AU111 | *Non identified* | Yeki | Shaka | Semi forest |
| AU112 | *T. viride* | Yeki | Shaka | Garden Coffee |
| AU113 | *T. asperellum* | Yeki | Shaka | Garden Coffee |

| | | | | |
|---|---|---|---|---|
| AU114 | *T. longibrachiatum* | Yeki | Shaka | Garden Coffee |
| AU115 | *T. asperellum* | Yeki | Shaka | Garden Coffee |
| AU116 | *T. citrinoviride* | Yeki | Shaka | Semi forest |
| AU117 | *Non identified* | Yeki | Shaka | Semi forest |
| AU118 | *T. asperelloides* | Limmu Saka | Jimma | Semi forest |
| AU119 | *T. asperelloides* | Yeki | Shaka | Semi forest |
| AU120 | *T. asperelloides* | Yeki | Shaka | Semi forest |
| AU121 | *T. longibrachiatum* | Sheko | Benchi Maji | Semi forest |
| AU122 | *T. asperelloides* | Yeki | Shaka | Semi forest |
| AU123 | *T. paratroviride* | Limmu Saka | Jimma | Semi forest |
| AU124 | *T. asperelloides* | Gera | Jimma | Semi forest |
| AU125 | *T. longibrachiatum* | Yayu | Buno Bedele | Forest |
| AU126 | *T. asperellum* | Yayu | Buno Bedele | Forest |
| AU127 | *Non identified* | Yayu | Buno Bedele | Forest |
| AU128 | *T. asperellum* | Yayu | Buno Bedele | Forest |
| AU129 | *T. asperellum* | Yayu | Buno Bedele | Forest |
| AU130 | *T. asperellum* | Uraga | West Guji | Garden Coffee |
| AU131 | *T. asperellum* | Gera | Jimma | Forest Coffee |
| AU132 | *T. hamatum* | Yirga cheffe | Gedeo | Garden Coffee |
| AU133 | *T. asperellum* | Sheko | Benchi Maji | Semi forest |
| AU134 | *T. asperelloides* | Sheko | Benchi Maji | Semi forest |
| AU135 | *T. asperelloides* | Sheko | Benchi Maji | Semi forest |
| AU136 | *T. longibrachiatum* | Sheko | Benchi Maji | Forest |
| AU137 | *T. asperellum* | Sheko | Benchi Maji | Forest |
| AU138 | *T. longibrachiatum* | Semien Benchi | Benchi Maji | semi forest |
| AU139 | *T. asperellum* | Semein Benchi | Benchi Maji | semi forest |
| AU140 | *T. longibrachiatum* | Semein Benchi | Benchi Maji | semi forest |
| AU141 | *T. longibrachiatum* | Semein Benchi | Benchi Maji | semi forest |
| AU142 | *T. asperellum* | Sheko | Benchi Maji | semi forest |
| AU143 | *T. longibrachiatum* | Sheko | Benchi Maji | semi forest |
| AU144 | *T. asperelloides* | Sheko | Benchi Maji | Forest |
| AU145 | *T. reesei* | Sheko | Benchi Maji | Forest |
| AU146 | *T. asperelloides* | Sheko | Benchi Maji | Forest |
| AU147 | *Non identified* | Sheko | Benchi Maji | Forest |
| AU148 | *T. asperelloides* | Haru | West Wollega | Forest |
| AU149 | *T. asperelloides* | Haru | West Wollega | Semi forest |
| AU150 | *T. gamsii* | Haru | West Wollega | Semi forest |
| AU151 | *Non identified* | Haru | West Wollega | Semi forest |
| AU152 | *T. longibrachiatum* | Limmu Saka | Jimma | Semi forest |
| AU153 | *T. asperellum* | Delo Mena | Bale | Forest |
| AU154 | *T. harzianum* | Delo Mena | Bale | Forest |
| AU155 | *T. asperelloides* | Delo Mena | Bale | Forest |
| AU156 | *T. asperelloides* | Delo Mena | Bale | Forest |
| AU157 | *T. aethiopicum* | Berbere | Bale | Forest |
| AU158 | *T. longibrachiatum* | Yeki | Sheka | Forest |
| AU159 | *T. brevicompactum* | Berbere | Bale | Forest |
| AU160 | *T. gamsii* | Berbere | Bale | Forest |
| AU161 | *T. longibrachiatum* | Berbere | Bale | Forest |
| AU162 | *T. asperelloides* | Kercha | West Guji | Garden Coffee |
| AU163 | *T. asperellum* | Jarso | West Wollega | Semi forest |
| AU164 | *T. longibrachiatum* | Jarso | West Wollega | Semi forest |

| | | | | |
|---|---|---|---|---|
| AU165 | *T. asperellum* | Aira Guliso | West Wollega | Semi forest |
| AU166 | *T. hamatum* | Semein Benchi | Benchi Maji | Semi forest |
| AU167 | *T. bissettii* | Bule Hora | West Guji | Garden Coffee |
| AU168 | *T. aethiopicum* | Kercha | West Guji | Garden Coffee |
| AU169 | *T. asperellum* | Kercha | West Guji | Garden Coffee |
| AU170 | *T. asperellum* | Aira Guliso | West Wollega | Garden Coffee |
| AU171 | *T. asperellum* | Aira Guliso | West Wollega | Garden Coffee |
| AU172 | *T. asperellum* | Kercha | West Guji | Garden Coffee |
| AU173 | *T. longibrachiatum* | Bule Hora | West Guji | Garden Coffee |
| AU174 | *T. asperellum* | Bule Hora | West Guji | Garden Coffee |
| AU175 | *T. asperelloides* | Bedele | Buno Bedele | Forest |
